# Capturing Snapshots of Nucleosomal H2A K13/K15 Ubiquitination Mediated by the Monomeric E3 Ligase RNF168

**DOI:** 10.1101/2024.01.02.573964

**Authors:** Huasong Ai, Zebin Tong, Zhiheng Deng, Qiang Shi, Shixian Tao, Jiawei Liang, Maoshen Sun, Xiangwei Wu, Qingyun Zheng, Lujun Liang, Jia-Bin Li, Shuai Gao, Changlin Tian, Lei Liu, Man Pan

## Abstract

The DNA damage repair regulatory protein RNF168, a monomeric RING-type E3 ligase, plays a crucial role in regulating cell fate and DNA repair by specific and efficient ubiquitination of the adjacent Lys13 and Lys15 sites at the H2A N-terminal tail. However, understanding how RNF168 coordinates with its cognate E2 enzyme UbcH5c to ubiquitinate H2AK13/15 site-specifically has long been hampered by the lack of high-resolution structures of RNF168 and UbcH5c∼Ub in complex with nucleosomes. Here, we developed mechanism-based chemical trapping strategies and determined the cryo-EM structures of the RNF168/UbcH5c–Ub/NCP complex captured in transient H2AK13/15 monoubiquitination and adjacent dual-monoubiquitination reactions. Our structural analysis revealed that RNF168 stably binds to the nucleosomal H2A–H2B acidic patch through a basic helix with multiple interactions, which positions the UbcH5c active centre directly over the H2A N-terminus, providing a “helix-anchoring” mode for monomeric E3 ligase RNF168 on nucleosome in contrast to the “compass-binding” mode of dimeric E3 ligases. Furthermore, our chemically synthesized ubiquitinated histones have enabled the elucidation of the efficiency of Ub installation and the interplay between the initial and subsequent Ub modifications on the adjacent H2A K13 and K15 sites. Overall, our work not only provides structural snapshots of H2A K13/K15 site-specific monoubiquitination and adjacent dual-monoubiquitination, but also offers a near-atomic resolution structural framework for understanding how pathogenic mutations or physiological modifications affect the molecular function of RNF168 in H2A K13/15 ubiquitination.

## Introduction

In response to DNA double-strand breaks, mammalian cells initiate a multi-component signaling cascade that involves various protein post-translational modifications such as phosphorylation, methylation, and ubiquitination^1–4^. A variety of DNA damage response proteins are recruited to the chromatin near the DNA breaks to orchestrate the proper damage repair and signaling. As a crucial chromatin-related factor for this process, the ubiquitin E3 ligase RNF168, which ubiquitinates histone H2A (or H2AX) at the N-terminal Lys 13/15 residues^5–7^, plays a pivotal role in the proper cellular response to DNA damage repair. The RNF168-driven H2A K13/15 ubiquitination provides a docking platform for the recruitment of a cohort of downstream DNA repair proteins at damaged chromatin sites to trigger the activation of distinct DNA repair pathways^8,9^, including the nonhomologous end-joining (NHEJ) pathway through proteins such as p53 binding protein 1 (53BP1)^10^ and the homologous recombination (HR) pathway through BRCA1/BARD1, RAD18, and RNF169^11–13^. The RNF168-mediated H2A K13/15 ubiquitination signalling was at the crossroads of the repair pathway selection of DNA double-strand breaks, which are toxic lesions that become more prevalent during tumorigenesis^14^. The aberrant RNF168 ubiquitin signalling pathway is intimately linked to chronic proteotoxic and genotoxic stress adaptations^15^. Dysfunction or mutations of RNF168 have been strongly implicated in the development and progression of various human diseases, including tumours such as breast and esophageal cancers^16^, as well as RIDDLE syndrome^17^, a novel human immunodeficiency and radiosensitivity disorder associated with defective DNA double-strand break repair. Consequently, mechanistic studies targeting RNF168 ubiquitination have garnered significant attention due to their profound biological and clinical implications^18,19^.

Since the discovery of histone H2A (or H2AX) ubiquitination by RNF168 in 2009^5^, progressive studies have been conducted to identify the substrate lysine site and understand how RNF168 achieves site-specific ubiquitination of H2A at K13/K15 on nucleosomes^6,7,20–22^. *In vivo* mass spectrometry and *in vitro* biochemical reconstitution investigations have established that RNF168 specifically mediates the monoubiquitination of H2A at both the K13 and K15 sites, as well as the dual-monoubiquitination at K13/K15^6,7,20^. A variety of biochemical and biophysical experiments, including NMR interaction mapping, enzymatic mutagenesis, cross-linking mass spectrometry, and data-driven modeling, have been performed to gain insights into the molecular mechanism of RNF168 action on nucleosome^20–22^. Nonetheless, full understanding of how RNF168 cooperates with its cognate E2 enzyme UbcH5c to perform site-specific ubiquitination of H2A K13/15 and simultaneous dual-monoubiquitination has long been hampered by the lack of high-resolution structures of RNF168/UbcH5c/nucleosomes complex, which would provide the atomic details of the interactions between RNF168, UbcH5c and nucleosomes.

Capturing of the trimeric RNF168/UbcH5c∼Ub complex on nucleosomes has been challenging due to the weak, transient, and dynamic interactions between RNF168, UbcH5c and nucleosomes, thus impeding the crystal or cryo-EM structure elucidation. Recently, chemical protein synthesis-based trapping strategies developed by our team^23–25^ and other research groups^26–33^ have proven to be feasible approaches for obtaining the stable complex of E3-E2-substrate complex. Here, we developed and utilized mechanism-based chemical trapping strategies by constructing stable ubiquitination intermediate mimics to determine the structures of the RNF168/UbcH5c∼Ub/nucleosome complex. These structures show that RNF168 binds the nucleosomal H2A–H2B acidic patch through a basic helix with multiple interactions, which positions the active center of UbcH5c directly over the H2A K13/15, providing a “helix-anchoring” mode for monomeric E3 ligase on nucleosome. Furthermore, the chemically synthesized ubiquitinated histones allow us to enzymatically investigate the efficiency of Ub installation and the interplay between the initial and subsequent ubiquitin modifications on the adjacent H2AK13 and K15. Our work gives the structural snapshots of H2AK13/15 site-specific ubiquitination and adjacent dual-monoubiquitination, revealing a unique “helix-anchoring” mode employed by monomeric E3 ligase on nucleosomes compared to dimeric E3 ligases. Additionally, our study provides a structural framework at near-atomic resolution for understanding the disease mutations in RNF168’s basic helix.

## Results

### RNF168 preferentially mediates dual-monoubiquitination of adjacent H2A K13 and K15 sites

We started our study with a biochemical reconstruction of RNF168 activity on the adjacent H2A K13 and K15 sites. RNF168-mediated nucleosomal ubiquitination has been reported to be specifically targeted to histone H2A K13 and K15^6,20^. RNF168^RING^ (residues 1-113) was reported to be the minimal construct that still retained H2A K13/K15 specificity (**Fig. 1a**)^20^. This construct (i.e., RNF168^RING^) was used in our subsequent biochemical and structural studies of RNF168. The strict H2A K13/K15 site specificity of RNF168 was confirmed by our biochemical experiment showing that RNF168 has little or no activity on H2A K13R/K15R-mutated NCP (**Fig. 1b**). To investigate whether there is a difference in RNF168 activity between the H2A K13 and K15 sites, the H2A K13/K15R and H2A K13R/K15 mutated NCPs were used as substrates for RNF168-mediated ubiquitination, and a slight and nonobvious preference of RNF168 for the H2A K13/K15R NCP was observed (**Fig. 1b**).

**Figure 1.**
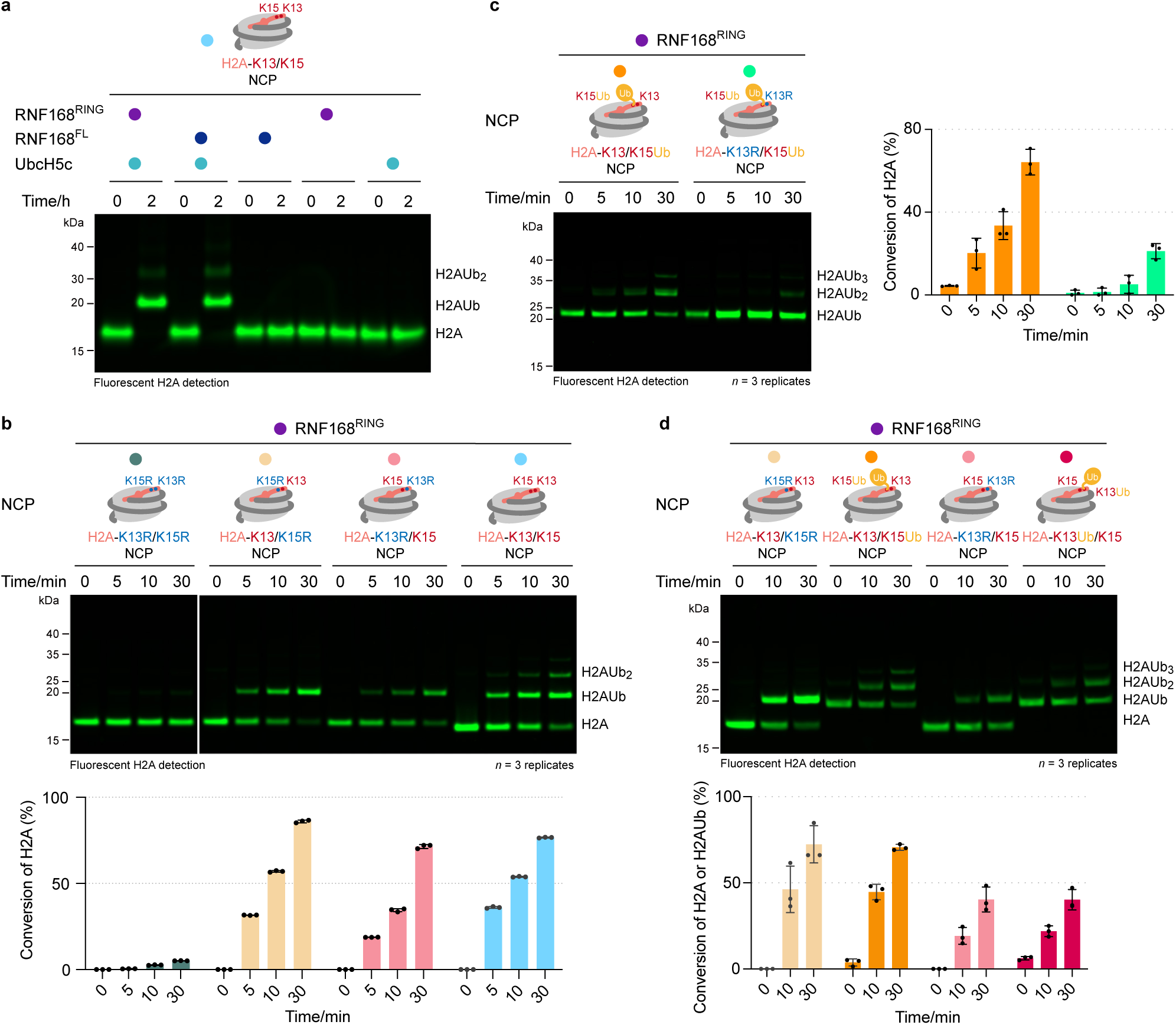
RNF168 efficiently mediates dual-monoubiquitination on adjacent H2A K13 and K15 sites. **a**, *In vitro* ubiquitination assay to compare the activity of RNF168^RING^ and RNF168^FL^ to generate monoubiquitination (H2AUb) and dual-monoubiquitination (H2AUb_2_) on H2A K13/K15 NCP. **b**, Top: *in vitro* ubiquitination assays to test the site-specificity of RNF168^RING^ on nucleosomal H2A K13 and K15 sites and to compare the ubiquitination efficiency of RNF168^RING^ on nucleosomal H2A K13 or K15 site using H2A K13/K15R NCP or H2A K13R/K15 NCP, respectively. Bottom: quantification of nucleosomal H2A ubiquitination in the top panel. **c**, Left: *in vitro* ubiquitination assay using H2A K13/K15Ub NCP and the corresponding K13R mutant (H2A K13R/K15Ub NCP) shows that RNF168^RING^ preferred to generate adjacent dual-ubiquitination on H2A K13 and K15 rather than mediate polyubiquitination chain elongation on the Ub motif of H2AUb. Right: quantification of nucleosomal H2A ubiquitination in the left panel. **d**, Top: *in vitro* ubiquitination assays to investigate the ubiquitination efficiency of RNF168^RING^ in the first monoubiquitination and the second ubiquitination steps of nucleosomal H2A K13/K15 adjacent dual-ubiquitination. The slight difference in band shift observed for H2AUb may be attributed to the presence of different native isopeptide linkage-mediate by RNF168 ubiquitination and the mimicked one-mediated by the CAET trifunctional molecule. Bottom: quantification of nucleosomal H2A ubiquitination in the top panel. All quantified nucleosomal H2A ubiquitination data in **Figure 1** show the value (black dots) and mean ± SD (bars) from *n* = 3 independent biological replicates. Notes: the scheme of the nucleosome disc was asymmetric but the symmetrically modified nucleosomes were used in biochemical experiments. The H2A variants in different nucleosome contexts used here in **Figure 1**, and in **Figures 4g, 4i** and **Figure 5f**, **Extended Data Figure 1**, **Extended Data Figures 13c**, **13h**, **13i**, and **13j**, and **Extended Data Figures 14** were derived from the H2A K0 variant in which all the lysines on H2A were substituted by arginines. For example, the H2A K13/K15 NCP used in **Figure 1a** indicates that only two lysines (K13 and K15) are present on H2A, while all other H2A lysines are replaced by arginines. Detailed sequences of H2A variants in different nucleosome contexts used in biochemical assays are listed in **Supplementary Table 3**.

H2A K13/K15 ubiquitination has been observed as polyubiquitination *in vivo*, with K27 and K63 ubiquitin linkages reported^17,34^. However, we only observed the appearance of shorter ubiquitin chain type on H2A, either using full-length RNF168 (residues 1–571), RNF168 (1–113) or a prolonged reaction time of 18 hours in our *in vitro* biochemical reconstitution. This phenomenon is also observed in previous studies^6,20–22^ (**Fig. 1a** and **Extended Data Fig. 1a–c**). Moreover, there was no difference in RNF168 ubiquitination activity when performing ubiquitination assays using wild-type (WT) or seven Lys-to-Arg mutated Ub (K6R, K11R, K27R, K29R, K33R, K48R, and K63R) as donor Ub on the H2A K13R/K15 NCP or chemically synthesized H2A K13R/K15Ub NCP (synthetic scheme and validation were shown in **Supplementary Fig. 1, 2**), suggesting that, at least under our *in vitro* experimental conditions, RNF168 does not have Ub chain-type specificity for H2A ubiquitination (**Extended Data Fig. 1c–d**). Conversely, we observed significant reduction of RNF168 activity in the H2A K13R/K15Ub NCP compared to the H2A K13/K15Ub NCP (**Fig. 1c**), indicating that when monoubiquitination of H2A occurs, the preferred ubiquitination event in the later stage is adjacent dual-ubiquitination, but polyubiquitination does not extend to the Ub motif of H2AUb, which is consistent with the findings of a previous study showing that multiple monoubiquitination of histone H2A occurred when a Ub variant without lysines (K0-Ub) was used^6^. This result also echoes the finding of the mass spectrometry analysis showing that both H2A K13 and K15 sites were co-modified with Ub *in vivo* in an RNF168-dependent manner^7^.

We then compared the ubiquitination efficiency in the first and second steps of H2A K13/K15 adjacent ubiquitination by using H2A K13/K15R NCP, H2A K13R/K15 NCP, chemically synthesized H2A K13Ub/K15 NCP, and H2A K13/K15Ub NCP (synthetic scheme in **Supplementary Fig. 1**). Similar ubiquitination efficiencies were observed in the same sites (K13 or K15), regardless of whether the adjacent site was modified with Ub or not (**Fig. 1d**), indicating that the pre-existing Ub at the H2A K13 or K15 sites does not act as a steric hindrance to the activity of RNF168.

### Structural capture of the RNF168/UbcH5c–Ub/NCP complex

An understanding of the structural mechanism of nucleosome-specific H2A K13/K15 ubiquitination has long been pursued but has been hampered by difficulties in obtaining stable RNF168/UbcH5c–Ub/NCP complexes. In our initial experiments, we also made several unsuccessful attempts, including 1) direct incubation of unmodified nucleosomes, full-length RNF168, and oxyester-bound UbcH5c∼Ub (**Extended Data Fig. 2a**) and 2) incubation of the unmodified nucleosomes with the full-length RNF168 (residues 1–571) or RNF168^RING^ that was linearly fused to UbcH5c (**Extended Data Fig. 2b-c**). Neither of these approaches was able to achieve a high-resolution cryo-EM reconstruction of RNF168, UbcH5c and the nucleosome complex, suggesting that their interactions may be weak and transient.

Thus, we turned to our recently developed intein-based E2-Ub-nucleosome conjugation strategy^24,25^ that allows the covalent junction of the E2 active centre, Ub C-terminus and nucleosomal substrate lysine residue (**Fig. 2a**). Briefly, the gp41-1 intein C-terminal fragment-fused E2 (referred to as IntC-E2) active centre, the Ub C-terminus, and the histone H2A K15^C^ (Lys15 was mutated into Cys) are chemically conjugated through a trifunctional molecule (CAET, 2-((2-chloroethyl)amino)ethane-1-(S-acetaminomethyl)thiol) (**Extended Data Fig. 3a**), and the synthesized IntC-E2-Ub-H2A is assembled into nucleosomes. Upon the addition of the gp41-1 intein N-terminal fragment-fused E3 (referred to as E3-IntN), protein splicing between the split IntC and IntN occurs, leading to the formation of covalent E3/E2–Ub/NCP conjugates (**Fig. 2a–b** and **Extended Data Fig. 3**). The stable ubiquitination transferring intermediate mimics (i.e., the E2–Ub–nucleosome conjugates) are of high similarity to the native trans-thioesterification state (**Fig. 2c–d**). Using this intein-based E2-Ub-nucleosome conjugation strategy, the RNF168/UbcH5c–Ub/NCP complex was prepared and subjected to single-particle cryo-EM analysis to obtain an overall 3.27 Å map of RNF168/UbcH5c–Ub/NCP (**Fig. 2e** and **Extended Data Fig. 4**), allowing us to unambiguously dock the structures of the RNF168 RING domain (PDB: 4GB0^35^), UbcH5c (PDB: 5EGG^36^), Ub (PDB: 1UBQ^37^) and nucleosome (PDB: 7XD1^38^) into the cryo-EM map and model the complex (**Fig. 2f** and **Extended Data Fig. 12a–h**).

**Figure 2.**
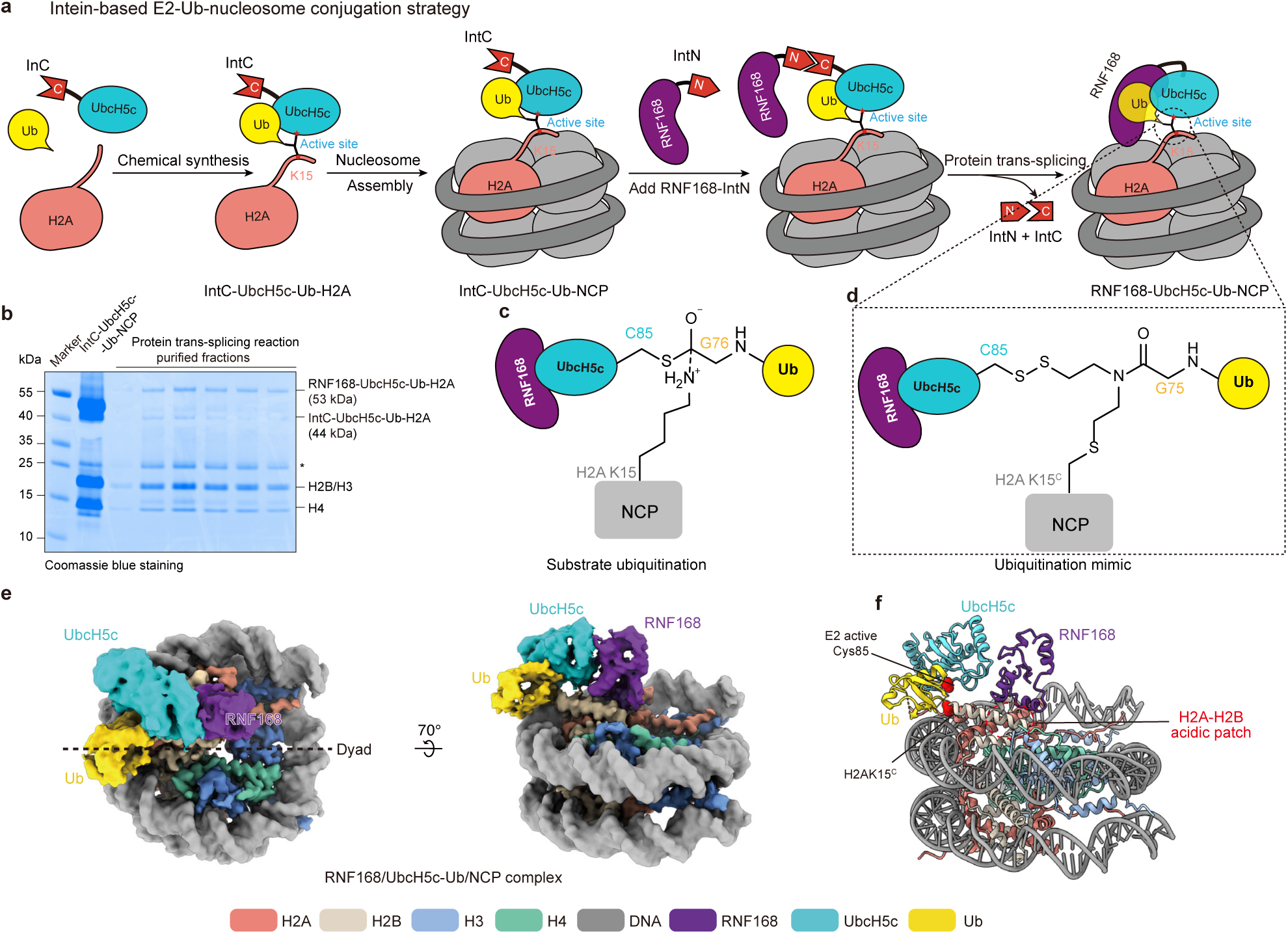
Capturing the complex of RNF168/UbcH5c-mediated nucleosome monoubiquitination by intein-based E2-Ub-nucleosome conjugation strategy. **a.** Schematic representation of intein-based E2–Ub–nucleosome conjugation strategy. The detailed synthetic route is shown in **Extended Data Figure 3a**. **b.** A coomassie-stained non-reducing SDS–PAGE gel of reconstituted IntC-UbcH5c-Ub-H2A nucleosome and purified RNF168-UbcH5c-Ub NCP. **c.** A schematic representation of the native transition state of nucleosomal H2A ubiquitination by RNF168-UbcH5c. RNF168 coordinates with UbcH5c-Ub to complete substrate ubiquitination in a tetrahedral transition state. **d.** The dashed box shows the designed intermediate structure by intein-based E2–Ub–nucleosome conjugation strategy, which highly mimics the transition state of the substrate ubiquitination by RNF168-UbcH5c. **e.** The 3.27 Å cryo-EM density map of RNF168/UbcH5c–Ub/NCP complex obtained by intein-based E2– Ub–nucleosome conjugation strategy. The map of the RNF168/UbcH5c–Ub/NCP complex was sharpened using a B factor of −30 Å^2^ and contoured at a level of 0.01. **f.** Atomic model of the RNF168/UbcH5c–Ub/NCP complex. The side chains of H2A K15 (mutated to cysteine) and UbcH5c (C85) are depicted as red spheres, respectively. Both are spatially adjacent.

In this reconstitution, the RNF168 RING domain is docked on the H2A–H2B acidic patch via a basic helix and binds to UbcH5c, positioning it towards the SHL DNA 4.5 and placing its active centre (Cys85) over the H2A K13/K15 sites (**Fig. 2e–f**). The placement of the Ub motif between the back of UbcH5c and the nucleosomal DNA at SHL 3.5 (**Fig. 2e–f**) is unexpected and differs from the typical positioning found in the canonical E3–E2 brace^39^. It should be noted that the manner in which Ub, H2A, and UbcH5c are conjugated through the CAET trifunctional molecule and the gp41-1 intein linkage between RNF168 and UbcH5c appear to have no effect on the overall conformational arrangement of RNF168, UbcH5c, and Ub on the nucleosome (**Extended Data Fig. 5** and **Supplementary Fig. 3**).

### Structural visualization of RNF168-mediated dual-monoubiquitination of the adjacent H2A K13 and H2A K15 sites

We additionally tried to delve into the structural mechanism of the subsequent RNF168 ubiquitination event on monoubiquitinated histone H2A, and attempted to chemically assemble the Ub C-terminus, UbcH5c, and the monoubiquitinated nucleosome substrate by using the same strategy described above (**Fig. 2a**). Considering that histone H2A K13Ub-K15^C^ is easier to synthesize than H2A K13^C^-K15Ub (where the K15C residue can serve as a ligation site for hydrazide-based native chemical ligation^40^), UbcH5c∼Ub was designed to be conjugated to histone H2A K13Ub-K15^C^. However, we failed to conjugate the chemically synthesized UbcH5c– Ub with histone H2A K13Ub-K15^C^ via disulfide bond exchange reactions (**Extended Data Fig. 6a**). This may be due to steric hindrance effects caused by the pre-existed Ub at histone H2A, making it challenging for the E2–Ub active centre to approach H2A K15^C^ spatially in the absence of the E3 ligase, thereby hindering the reaction.

To overcome this problem, we were inspired by the ability of RNF168 to ubiquitinate on monoubiquitinated nucleosomes (**Fig. 1c-d**), and developed a novel activity-based chemical trapping strategy that exploits the E3 ligase mediated spatial proximity of the E2 enzyme and substrate site (as shown in **Fig. 3a**). Specifically, the IntC-E2 enzyme active centre and Ub C-terminus were chemically conjugated through a trifunctional molecule (CAET) to give IntC-E2-Ub, followed by the addition of E3-IntN to give the auto-spliced E3–E2–Ub complex. Meanwhile, the sulfhydryl group of the chemically synthesized H2A K13Ub-K15^C^ NCP was activated using 2,2′-dipyridyl disulfide (AT_2_) to afford the H2A K13Ub-K15^C-AT^ NCP (**Extended Data Fig. 6b-c**). Finally, the E3–E2–Ub complex was incubated with the H2A K13Ub-K15^C-AT^ NCP, and the sulfhydryl group in the E2–Ub active centre was covalently captured by the disulfide-activated H2A K13Ub-K15^C-AT^ NCP (**Fig. 3b**), a process induced by enzyme-substrate proximity, mimicking the native substrate ubiquitination intermediates (**Fig. 3c-d**).

**Figure 3.**
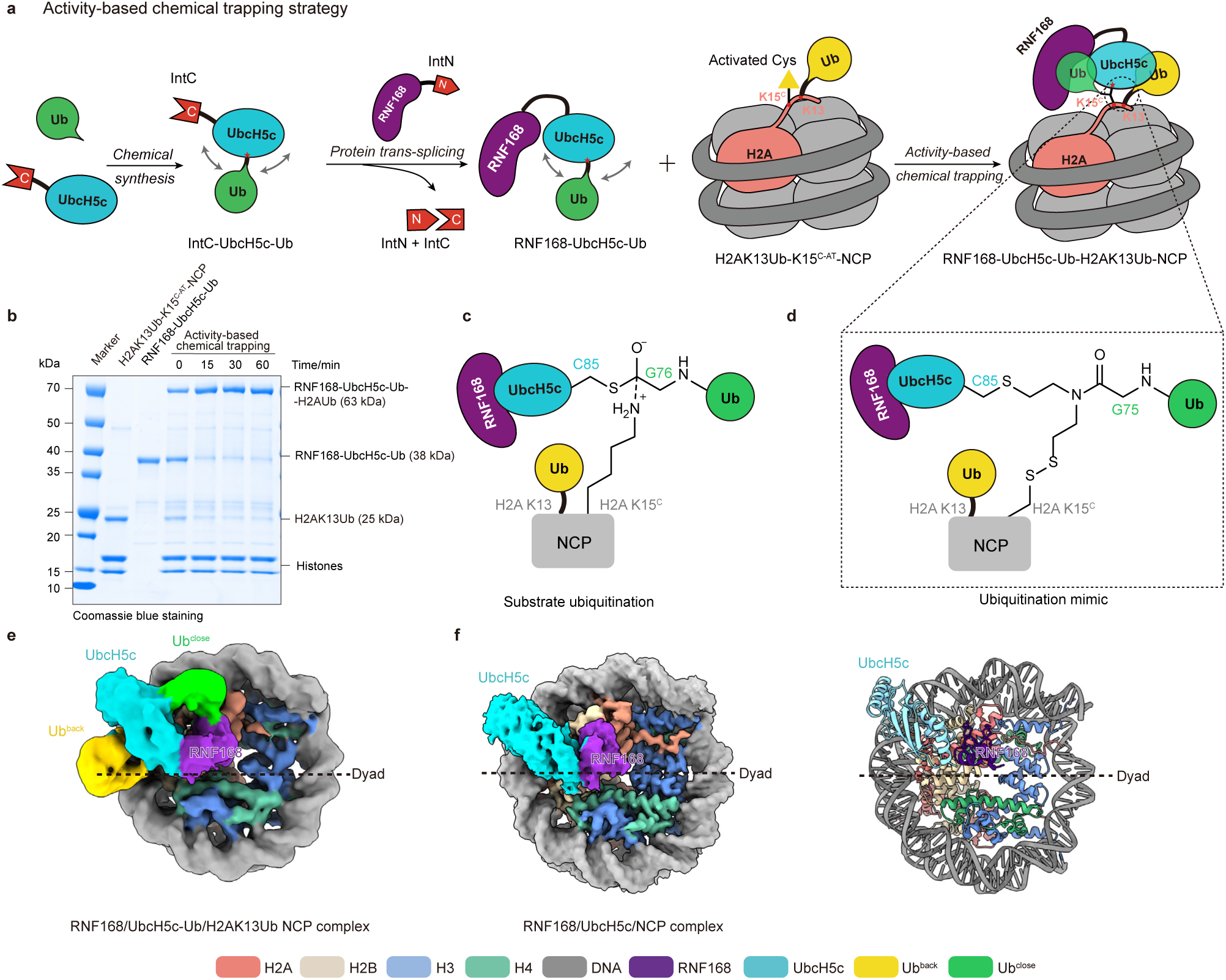
Structural Visualization of RNF168/UbcH5c-mediated H2A K13/K15 dual-monoubiquitination by activity-based chemical trapping strategy. **a.** Schematic representation of the activity-based chemical trapping strategy used to capture RNF168-mediated adjacent dual-monoubiquitination. The detailed synthetic route is shown in **Extended Data Figure 6b**. **b.** A coomassie-stained non-reducing SDS–PAGE gel of reconstituted H2A K13Ub-K15^C-AT^ nucleosome, purified RNF168-UbcH5c-Ub, and aliquots of activity-based chemical trapping reaction using RNF168-UbcH5c-Ub and H2A K15^C-AT^ nucleosome taken at 0.1 min,15 min, 30 min, 60 min. According to the molecular weight from the largest to the smallest, the bands on the gel are according to the following, respectively: RNF168-UbcH5c-Ub-H2A K13Ub (63 kDa), RNF168-UbcH5c-Ub (38 kDa), H2A K13UbK15^C-AT^ (25 kDa), H3 (15 kDa), H2B (13 kDa) H4 (11 kDa). **c.** A schematic representation of the native transition state of nucleosomal H2A dual-monoubiquitination at adjacent sites of H2A K13 and H2A K15 by RNF168-UbcH5c. **d.** The dashed box shows the designed intermediate structure by activity-based chemical trapping strategy, which highly mimics the transition state of the dual-monoubiquitination by RNF168-UbcH5c. **e.** The 6.25 Å cryo-EM density map of RNF168/UbcH5c∼Ub/H2A K13Ub-K15^C^ nucleosome complex obtained by activity-based chemical trapping strategy. The map of the complex was contoured at a level of 0.01. In this conformation, the density of the two Ub is visible. **f.** (Left) The 3.2 Å cryo-EM density map of RNF168/UbcH5c/NCP complex. In this conformation, the density of the two Ub is invisible. The map of the complex was sharpened using a B factor of −30 Å^2^ and contoured at a level of 0.0125. (Right) Atomic model of the RNF168/UbcH5c/NCP complex.

Using this activity-based chemical trapping strategy, the RNF168/UbcH5c– Ub/H2A K13Ub NCP complex was successfully obtained and subjected to cryo-EM analysis. A total of 8,480 micrographs were collected and processed with RELION 3.1 (**Extended Data Fig. 7**) to give a reconstruction of RNF168/UbcH5c/NCP complex at 3.2 Å (the Ub density was invisible) and a reconstruction of RNF168/UbcH5c–Ub/H2A K13Ub NCP complex at 6.25 Å (two Ub motifs were visualized, albeit at a low resolution) (**Fig. 3e–f** and **Extended Data Fig. 12i–o**). The overall conformation of RNF168 and UbcH5c on the nucleosome was similar in the two maps. In the 6.25 Å cryo-EM map, our ability to discern the fine details of ubiquitin interactions is restricted due to the low resolution, and our analysis is therefore primarily focused on identifying the spatial positioning of ubiquitin. Within this map, we observe two distinct Ub motifs: one Ub is located within the brace formed by RNF168 and UbcH5c, and this placement resembles the canonical closed E2∼Ub conformation (**Fig. 3e**). Consistent with this structure, the introduction of mutations in the conserved UbcH5c residues (I88A, L97A and L104A), which are critical for the formation of the closed E2∼Ub conformation, resulted in a substantial decrease in the RNF168 nucleosomal ubiquitination activity (**Extended Data Fig. 8a-b**). The other Ub motif was found to be situated between UbcH5c and SHL DNA 3.5, echoing a similar conformation captured in the RNF168-mediated initial monoubiquitination at H2A K15 (**Fig. 2e**). The presence of two spatially separated Ub conformations, one located within the brace formed by RNF168– UbcH5c and the other outside of it, aligns with the biochemical results (as shown in **Fig. 1d**) showing that the pre-existing Ub at the H2A K13 or K15 sites does not act as a steric hindrance to the RNF168-mediated dual-monoubiquitination of the adjacent H2A K13 and H2A K15 sites.

### Structural basis of monomeric E3 ligase RNF168 on the H2A–H2B acidic patch

In our two resolved complex structures of the RNF168/UbcH5c–Ub/NCP complex by the E2–Ub–nucleosome conjugation strategy (**Fig. 2e**) and the RNF168/UbcH5c– Ub/H2A K13Ub NCP complex by the activity-based chemical trapping strategy (**Fig. 3f**), the overall conformations and interaction patterns of RNF168 and UbcH5c on nucleosomes were highly similar, with a root-mean-square deviation (RMSD) of 0.337 Å when these structures are aligned using the nucleosome as a reference (**Fig. 4a-c**). To simplify analysis, we next performed a unified analysis of the detailed interactions among RNF168, UbcH5c, and nucleosomes using the structure determined by the E2– Ub–nucleosome conjugation strategy, which provided a higher local resolution. The results of this analysis are presented in the following sections.

**Figure 4.**
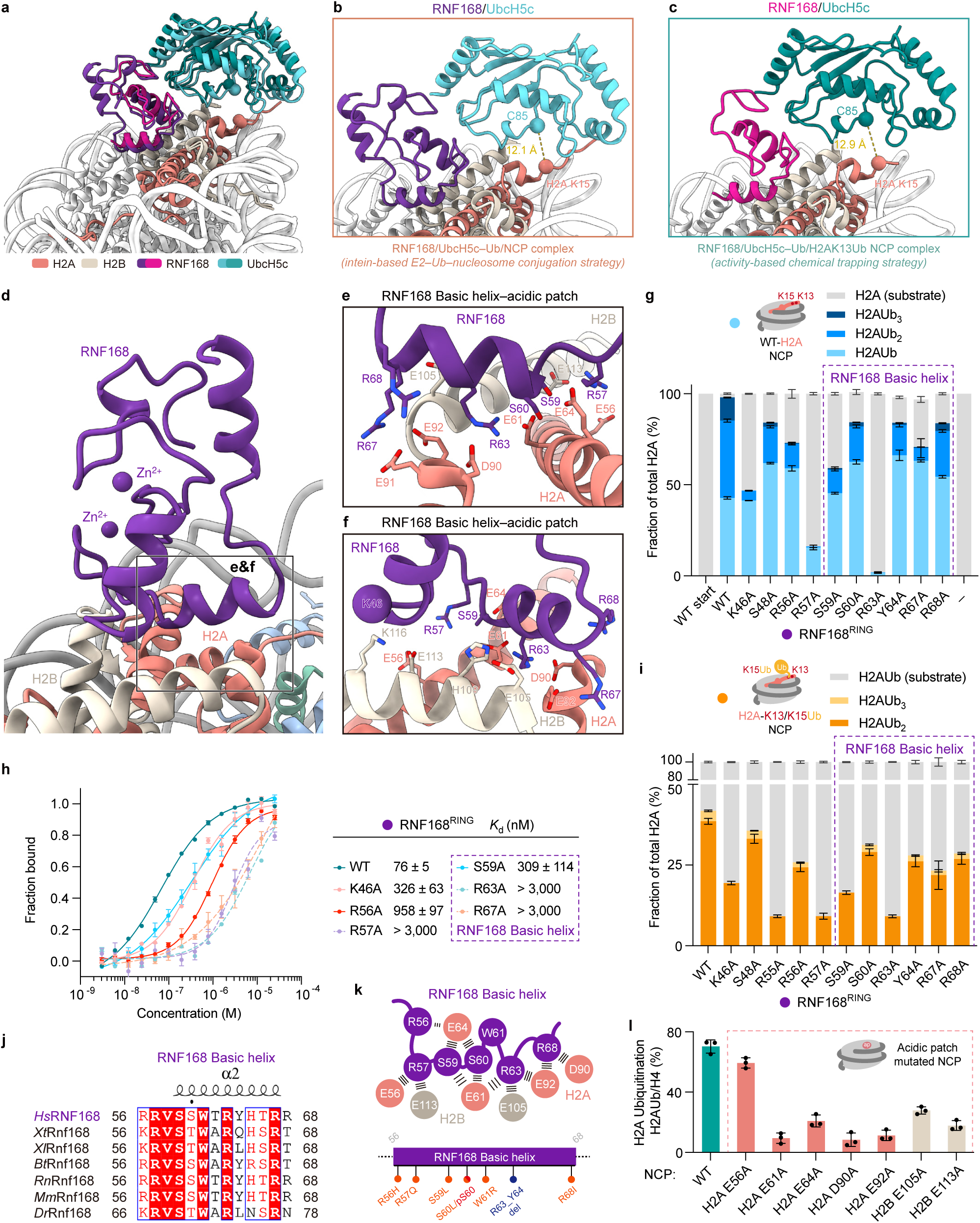
Structural basis of monomeric E3 ligase RNF168 on the H2A–H2B acidic patch. **a**, Structural alignment of RNF168/UbcH5c–Ub/NCP complex achieved by intein-based E2–Ub–nucleosome conjugation strategy, and the structure of RNF168/UbcH5c–Ub/H2A K13Ub NCP complex captured by activity-based chemical trapping strategy, highlighting the congruence in nucleosome-binding mode and E3 /E2 arrangement. **b**–**c**, Close-up views of RNF168/UbcH5c on the nucleosomal acidic patch in the two structures including the RNF168/UbcH5c–Ub/NCP complex by intein-based E2–Ub–nucleosome conjugation strategy (**b**), and the RNF168/UbcH5c–Ub/H2A K13Ub NCP complex by activity-based chemical trapping strategy **(c**). The Cαs of UbcH5c active center C85 and H2A K15 are depicted as spheres and the distances between them are indicated. **d**, Overview of interactions of RNF168^RING^ with the H2A– H2B acidic patch. The rectangle region indicates the close-up interfaces shown in **e** and **f**. The models shown in **d**–**f** are from the same model of the RNF168/UbcH5c–Ub/NCP complex achieved by intein-based E2–Ub–nucleosome conjugation strategy in **Figure 2f. e** and **f**, Different close-up views of the RNF168^RING^ basic helix–acidic patch interface with key interacting side chains shown. The alpha-carbon of RNF168 K46 residue was shown. **g**, Quantified nucleosomal H2A ubiquitination from *in vitro* ubiquitination assays using WT-H2A NCP to investigate the effects of RNF168^RING^ mutants on overall nucleosomal H2A ubiquitination pattern. **h**, Fluorescence polarization binding curves of WT RNF168^RING^ and mutants with fluorescently labeled NCP. Determined *K*_d_ values are indicated. Curves are fitted using data points from *n* = 3 independent biological replicates. The mean ± SD of each data point is indicated with bars. **i**, Quantified nucleosomal H2A ubiquitination from *in vitro* ubiquitination assays using H2A K13/K15Ub NCP to investigate the effects of RNF168^RING^ mutants on the second ubiquitination step of nucleosomal H2A K13/K15 adjacent dual-ubiquitination. **j**, Sequence alignment of RNF168 basic helix across species, highlighting the conservation of the basic helix. *Hs*, *Homo sapiens*; *Xt*, *Xenopus tropicalis*; *Xl*, *Xenopus laevis*; *Bt*, *Bos taurus*; *Rn*, *Rattus norvegicus*; *Mm*, *Mus musculus*; *Dr*, *Danio rerio*. *Hs*RNF168 secondary structure is depicted above the sequences. **k**, Top: Schematic RNF168^RING^ (residues 56-68) interface with key interacting residues shown. Bottom: A schematic representation for cancer mutations and post-translational modifications on RNF168 basic helix, reported in the Catalogue of Somatic Mutations in Cancer (COSMIC) database and reference^41^. **l**, Quantified nucleosomal H2A ubiquitination from *in vitro* ubiquitination assays using H2A–H2B acidic patch-mutated NCPs. Data in **g**, **i,** and **l** show the mean ± SD (bars) from *n* = 3 independent biological replicates.

An analysis of the interfaces between RNF168 and nucleosomes revealed that a basic helix of RNF168 (residues 59–68) lies on the H2A–H2B acidic patch formed by H2A residues (E61, E64, D90, E92) and H2B residues (E105, H109, E113), burying almost 180 Å^2^ of the solvent-accessible surface (**Fig. 4d–f**). The RNF168 R57 sidechain preceding the basic helix and R63 in the middle of the basic helix inserts into the deep acidic patch with charge interactions. The side chains of RNF168 residues S59, S60, and R67 in the helix are positioned towards the acidic patch (**Fig. 4d–f**). Additionally, the sidechain of the H2B residue K116 in the C-terminal helix is directed to the main chain of the RNF168 loop (E45/K46/A47) (**Fig. 4f**). These multiple interactions of the RNF168 RING domain on nucleosomes contribute to the stabilized conformation of the monomeric RNF168 E3 ligase on nucleosome surfaces.

In our biochemical validation experiments, mutations of the RNF168 residues to alanine (R56A, R57A, S59A, R63A, or R67A) led to a decrease in the nucleosomal ubiquitination activity on unmodified nucleosomes (**Fig. 4g**), as well as a weakening of the binding affinity with nucleosomes (**Fig. 4h**). Additionally, these mutations also resulted in a decrease in nucleosomal dual-monoubiquitination activity when using the monoubiquitinated nucleosome as a substrate (**Fig. 4i**), which provides additional biochemical evidence to support the structural observation that RNF168 exhibits very similar conformational binding to both unmodified and monoubiquitinated nucleosomes (**Fig. 4a-c**). A sequence alignment analysis of the RNF168 (residues 56-68) across various species, including *homo sapiens*, *Xenopus tropicalis*, *Xenopus laevis*, *Bos taurus*, *Rattus norvegicus*, *Mus musculus*, and *Danio rerio*, demonstrated the evolutionary conservation of their roles (**Fig. 4j**). Furthermore, introducing mutations in the nucleosomal H2A–H2B acidic patch, including H2A (E61A, E64A, D90A, E92A) or H2B (E105A, E115A), was found to significantly reduce RNF168-mediated nucleosome ubiquitination activity (**Fig. 4l**). The atomic-level structure we generated, combined with mutation analysis further refines the findings of previous biochemical investigations, confirming the crucial role of RNF168’s basic helix and nucleosomal H2A–H2B acidic patch in H2A K13/K15 ubiquitination^20,21^.

The phosphorylation of RNF168 at S60 by ribosomal S6 kinase has been reported to inhibit RNF168 function and hinder DNA damage repair^41^. Our structures offer a compelling explanation for this finding: residue S60 in the RNF168 basic helix is situated on the H2A–H2B acidic patch, with interactions with H2A E61 and E64 residues (**Fig. 4e-f**). The phosphorylation at residue S60, which introduces negatively charged groups, is likely to decrease the affinity of RNF168 for the nucleosome due to the repulsive forces generated by the electrical charges. Moreover, missense mutations, including R55C, R56H, R57Q, S59L, S60L, W61R, T68I point mutants^42–47^ in the RNF168 basic helix of RNF168, and the R63_Y64del deletion mutation^48^, have been identified in various cancers (**Fig. 4k** and **Supplementary Table 2**). For example, the RNF168 arginine anchor residue R57, which is crucial for nucleosomal H2A K13/K15 ubiquitination (**Fig. 4h–i**), has been found to be mutated into asparagine in sporadic intrahepatic cholangiocarcinoma^42^. Our structural elucidation of the RNF168 basic helix on the H2A–H2B acidic patch offers valuable insights into the functional implications of these mutations and contributes to our understanding of their involvement in disease development.

### Conserved E3-E2 interactions of RNF168 and UbcH5c

We aligned the interfaces among different nucleosomal E3 ligases (Ring1B/Bmi1^49,50^, BRCA1/BARD1^24,51,52^, Bre1^25^, and RNF20/40^25,53^) and their cognate E2 conjugation enzymes (UbcH5c, Rad6 or RAD6A) with the E2 enzyme as the aligning reference, and an analysis of the interfaces between the E2 enzymes and E2-binding RING domain of E3 ligases revealed a conserved E3–E2 interacting pattern^54^ (**Extended Data Fig. 8c**). The binding interface between RNF168 and UbcH5c is conventional and centred around the UbcH5c S94-P95-A96 motif (**Fig. 5a–b**). This motif is situated in proximity to RNF168 specific residues, including RNF168 loop residues P52/F53, I18 within the first Zn^2+^-binding loop, and T43 in the central helix of the RING motif (**Fig. 5a–b**). The introduction of the S94A mutation in UbcH5c leads to a reduction of approximately 75% in RNF168 ubiquitination activity (**Extended Data Fig. 8b**). Additionally, the UbcH5c residues S91 and N92, positioned after the helix (87–89), interact with RNF168 residue S48 and the linchpin residue R55, respectively (**Fig. 5c**). The introduction of RNF168 S48A or R55A leads to a decrease in the ubiquitination activity of RNF168 (**Fig. 4g, 4i**).

**Figure 5.**
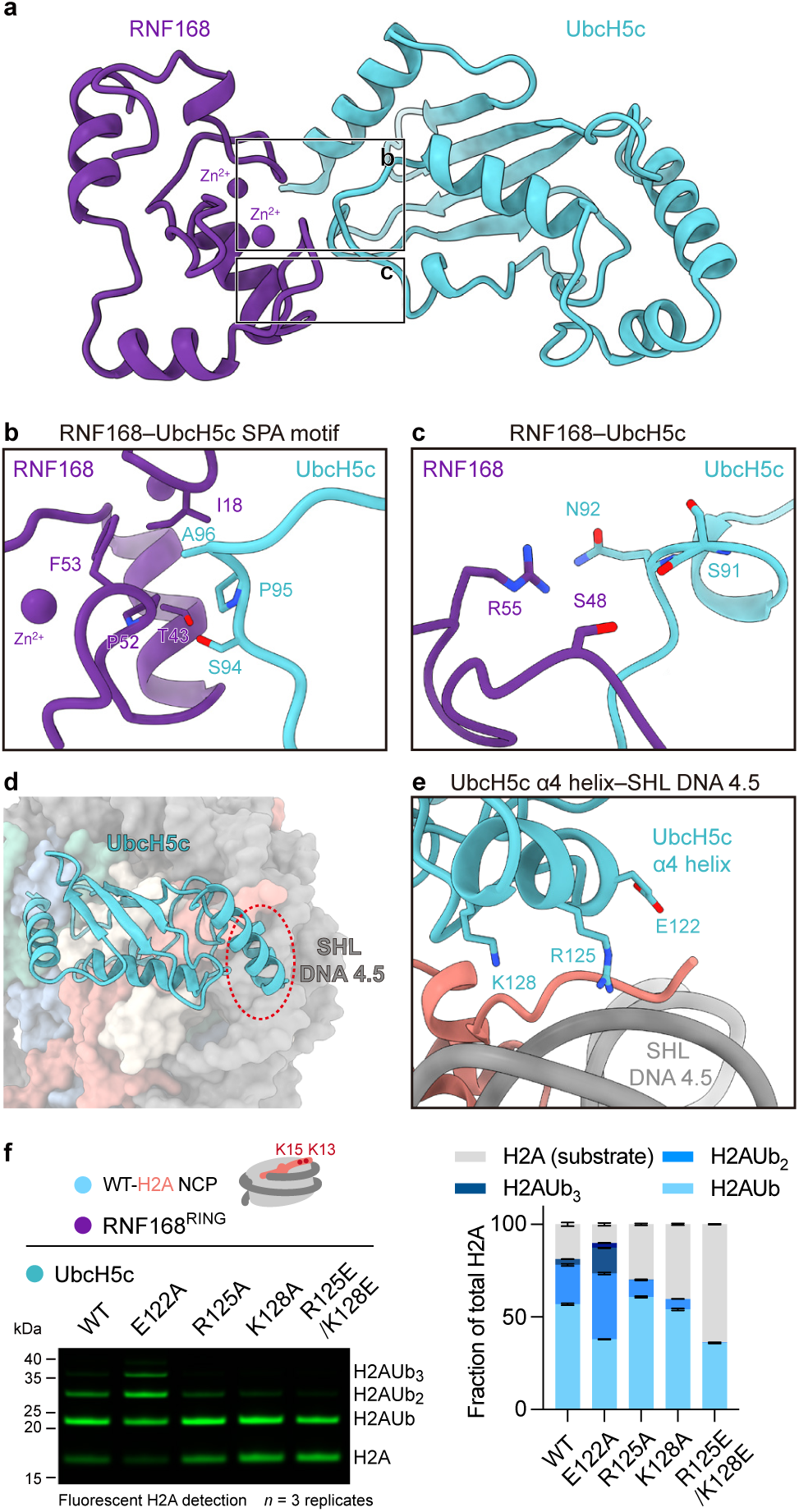
Analysis of interactions between UbcH5c with RNF168 and nucleosomal SHL DNA 4.5. **a**, Overview of interactions of RNF168^RING^ and UbcH5c. Rectangle regions indicate closed-up interfaces shown in **b** and **c**. The models shown in **a**–**e** are from the same model of the RNF168/UbcH5c–Ub/NCP complex achieved by intein-based E2–Ub–nucleosome conjugation strategy in **Figure 2f. b**, Close-up view highlighting the RNF168^RING^–UbcH5c SPA motif interface. **c**, Close-up view highlighting the interface centered on the RNF168^RING^ linchpin R55 and UbcH5c N92.**d**, Overview of UbcH5c on the nucleosome disk surface. The red dash oval circle highlights the interactions of the UbcH5c α4 helix with the SHL DNA 4.5. **e**, Close-up view showing the UbcH5c α4 helix–SHL DNA 4.5 interface. The side chains of UbcH5c E122, R125, and K128 are modeled based on the UbcH5c crystal structure (PDB: 5EGG). **f**, Left: *in vitro* ubiquitination assay using WT-H2A NCP to investigate the UbcH5c α4 helix–SHL DNA 4.5 interactions using WT UbcH5c and indicated mutants. Right: quantification of nucleosomal H2A ubiquitination in the left panel. Data show the mean ± SD (bars) from *n* = 3 independent biological replicates. Detailed sequences of WT H2A NCP used in biochemical assays are listed in **Supplementary Table 3**.

### Interactions of UbcH5c and nucleosomal DNA

With the fixed position of the RNF168 RING domain on nucleosomal H2A–H2B acidic patch, RNF168 interacts with UbcH5c through a canonical E3–E2 interface to position UbcH5c towards the SHL DNA 4.5, with the catalytic centre Cys85 directly over the H2A N-terminal tail near the K13/K15 sites (**Fig. 4a-c**). However, UbcH5c barely makes contact with the nucleosomal histone surface, and the lack of specific interactions between UbcH5c and the substrate may allow UbcH5c to efficiently ubiquitinate multiple substrates by pairing with different E3 ligases (e.g., RNF168^6^, Ring1B/Bmi1^55^ and BRCA1/BARD1^56^). Previous studies have suggested that the histone H2A α1 extension helix (16–22) plays a critical role in RNF168 activity^22^, whereas we did not observe direct engagement of UbcH5c with this helix in our structures.

Two basic residues (R125 and K128) of UbcH5c in the helix (121–129) were observed to be located above SHL DNA 4.5 (**Fig. 5d**). Although the density resolution of the helix prevents the modelling of sidechains, docking the sidechains of the two residues from the UbcH5c crystal structure (PDB: 5EGG)^36^ suggests potential for charge-interactions between R125/K128 and DNA (**Fig. 5e**). Mutations in UbcH5c, specifically R125A or K128A, resulted in a reduction in RNF168 ubiquitination activity, and the charge-reversal double mutation of R125E&K128E further reduced the activity (**Fig. 5f**). The UbcH5c residues R125 and K128 have also been previously reported to be involved in interactions with the nucleosomal dyad (SHL DNA 0) in structural studies of PRC1 Ring1B/Bmi1-mediated nucleosomal H2A K119 ubiquitination^49^. Our work thus provides another structural example demonstrating the essential involvement of E2 enzymes in recognizing nucleosomal DNA during nucleosome ubiquitination. We also noticed a negatively charged residue E122 in the helix located above the nucleosomal DNA; interestingly, a mutation abolishing the charge of UbcH5c E122 (E122A) stimulates RNF168-mediated nucleosomal ubiquitination (**Fig. 5f**).

### H2A K13/K15 specificity by RNF168

After analysing the detailed interactions of RNF168, UbcH5c and nucleosome, we turned to explore the nucleosomal H2A K13/15 specificity by RNF168. Previous near-atomic resolution structures of E3/E2/NCP complexes involving heterodimeric Ring1B/Bmi1^24,49^, BRCA1/BARD1^24,51,52^, RNF20/40^25^, and homodimeric Bre1^25,53^ have revealed a “compass-binding” mode of dimeric E3 ligases on the nucleosome (**Fig. 6a**). In the “compass-binding” mode, the E2-bound RING domain inserts into the H2A– H2B acidic patch via basic loop residues (K97/R98 in Ring1B, K70/R71 in BRCA1, R953/R955 in RNF20, and R679/R681 in Bre1A), while the non-E2 binding RING domain interacts with distinct nucleosomal regions, including the H3 α1–L1 elbow (Bmi1), the H2B–H4 cleft or H2B C-terminal helix (BARD1) and the nucleosomal SHL DNA 6.7 (RNF40 and Bre1)^25,53^. Instead of relying on interactions with multiple nucleosome spots, we found that RNF168 utilizes a basic helix (residues 59–68) to establish multiple interactions with the H2A–H2B acidic patch, thereby stabilizing the RNF168 RING domain on the nucleosome surface (**Fig. 6b**). This “helix-anchoring” mode employed by RNF168 to stabilize a monomeric E3 ligase on nucleosomes contrasts with the “compass-binding” mode of dimeric E3 ligases. The unique mode of binding exhibited by RNF168 allows it to effectively target nucleosomes without the need for dimerization. Through structurally conserved E3–E2 interactions, RNF168 precisely positions its corresponding E2 enzyme UbcH5c directly over the H2A N-terminus, with its active centre in close proximity to H2A K13/K15 (**Fig. 6b**).

**Figure 6.**
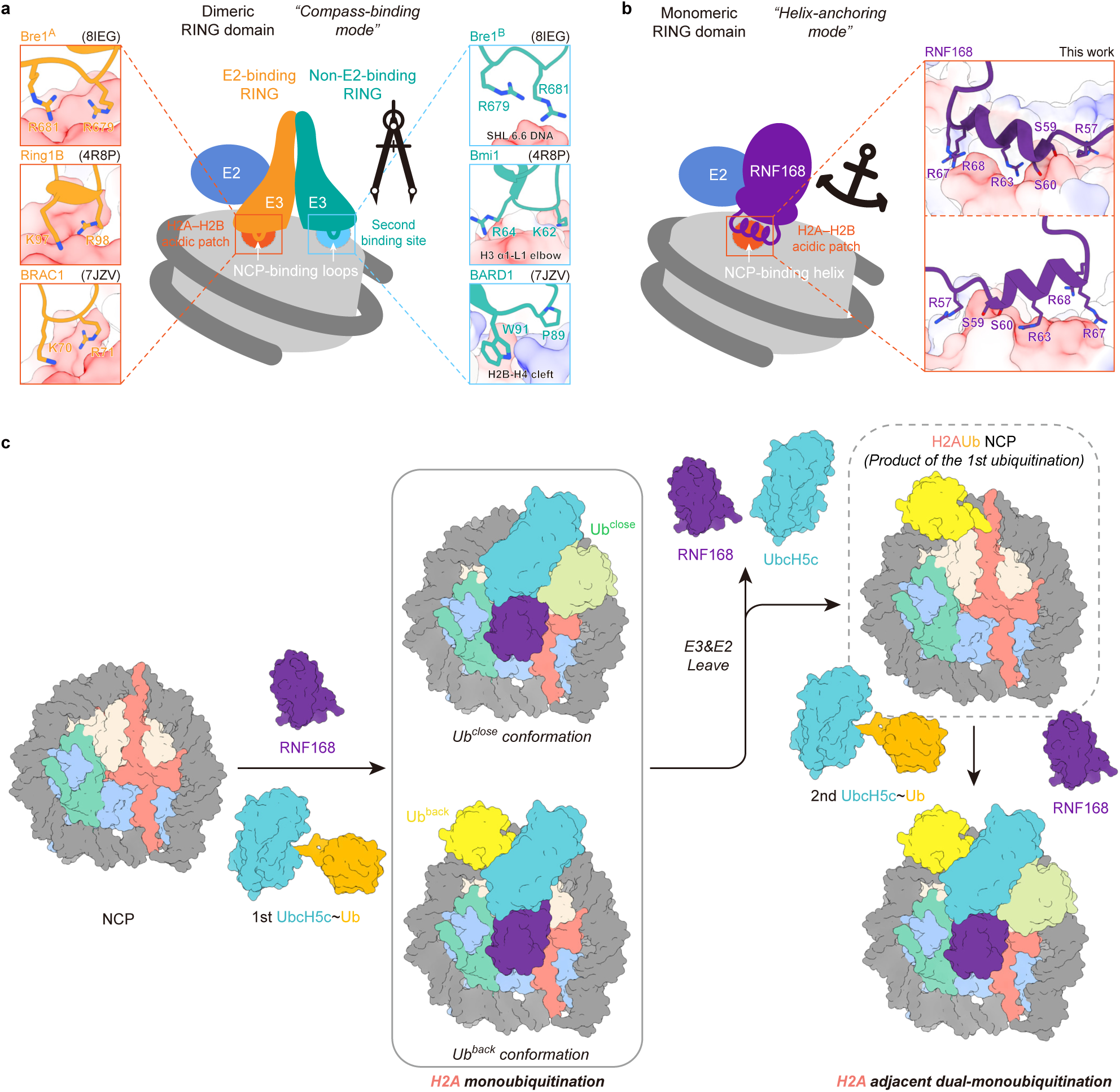
Model for nucleosomal H2A ubiquitination mediated by monomeric E3 ligase RNF168. **a**, A cartoon model illustrating how dimeric RING-type E3 ligases (including Bre1, Ring1/Bmi1, and BRAC1/BARD1) achieve site-specific ubiquitination on the nucleosome. Detailed interactions of the NCP-binding loops of the E2-binding RING domain and the non-E2-binding RING domain with the H2A– H2B acidic patch and the second binding sites, respectively, in the structures of Bre1 (PDB: 8IEG), Ring1/Bmi1 (PDB: 4R8P), and BRAC1/BARD1 (7JZV) are displayed. E3 ligases are shown in ribbon models and the nucleosome is depicted in an electrostatic potential surface. **b**, A cartoon model illustrating how the monomeric RING-type E3 ligase RNF168 catalyzes site-specific ubiquitination on the nucleosome. Detailed interactions of the NCP-binding helix of RNF168 with the H2A–H2B acidic patch are displayed. RNF168 is shown in a ribbon model and the nucleosome is depicted in an electrostatic potential surface. **c**, A cartoon model describing the mechanisms of RNF168-mediated H2A K13/K15 dual-monoubiquitination, including the first monoubiquitination step and the second adjacent monoubiquitination step.

Interestingly, RNF168 has been reported to exhibit specificity for H2A K13/K15 in the context of both the H2A–H2B dimer and the nucleosome^20^, distinguishing it from other nucleosomal E3 ligases, such as Ring1B/Bmi1^24,49^, BRCA1/BARD1^24,51,52^, RNF20/40^25^ and Bre1^25,53^, which are exclusively specific to nucleosome substrates. In our resolved RNF168/UbcH5c/NCP complex structures, RNF168 and UbcH5c exclusively interacted with histone H2A and H2B, without any observed interactions with histone H3 and H4 (**Fig. 4**). While other dimeric E3 ligases^24,25,49,51–53^ make contact not only with histone H2A and H2B, but also with spots involving histone H3, H4, or nucleosome-context dependent SHL DNA 6.7. This provides a plausible mechanistic basis for the RNF168 specificity for H2A–H2B dimers and nucleosomes. The potential interactions between UbcH5c and SHL DNA 4.5 (**Fig. 5e**) might be attributed to the slightly different ubiquitination activity previously observed for of RNF168 on H2A– H2B dimer and nucleosome substrates^20^.

## Discussion

When DNA damage occurs, RNF168 is recruited to the site of damage, playing a crucial role in initiating and coordinating the DNA damage response, ensuring efficient and accurate repair of DNA lesions and maintaining genomic integrity, all of which are activity that depends on RNF168’s ubiquitination activity on histone H2A lysine 13 and 15^1,18^. The structural mechanism underlying the site-specific ubiquitination by RNF168 has been studied over the past 10 years. The structural capture of the trimeric RNF168/UbcH5c∼Ub complex on nucleosomes has been a long challenge due to the weak, transient, and dynamic interactions between RNF168, UbcH5c, and nucleosomes, which has hindered the crystal or cryo-EM structure elucidation. Recently, chemical protein synthesis-based trapping strategies developed by our team^23–25^ and others^26–33^ have proven to be feasible approaches for obtaining the stable complex of E3-E2-substrate complex. Here, we first captured a snapshot of the RNF168/UbcH5c–Ub/NCP complex in the monoubiquitination reaction via our previously reported E2–Ub– nucleosome strategy^24,25^. Then, to circumvent the difficulty in the synthesis of E2-Ub-conjugated monoubiquitinated nucleosomes, we developed a novel activity-based chemical trapping strategy that exploits the E3 ligase-mediated spatial proximity of the E2 enzyme and substrate site to obtain a stable intermediate mimic, which captures the structures of RNF168-mediated adjacent dual-monoubiquitination of H2A K13/K15. The two complex structures reveal a congruent conformation and interaction details for the RNF168–UbcH5c ubiquitinating module on nucleosomes, providing the first structural snapshots of H2A K13/K15 site-specific ubiquitination at near-atomic resolution.

Notably, we also tried to use the activity-based chemical trapping strategy to study the RNF168-mediated monoubiquitination reaction (**Extended Data Fig. 9-11**). Through this approach, we successfully obtained a stable RNF168/UbcH5c–Ub/NCP complex with a cryo-EM reconstruction at 3.27 Å resolution (**Extended Data Fig. 10-11** and **Extended Data Fig. 12p-w**) and the structure of this complex closely resembled that determined using the E2–Ub–nucleosome conjugation strategy (**Fig. 2e**). Nonetheless, during the processing of this dataset, we identified a low-resolution (6.87 Å) conformation (**Extended Data Fig. 10d**) that was not observed in the complex captured through the E2–Ub–nucleosome conjugation strategy. In this low-resolution conformation, Ub was located in the braces of RNF168 and UbcH5c (**Extended Data Fig. 11f-g**), representing the closed E2∼Ub conformation. This conformation is consistent with our biochemical results showing that the closed E2∼Ub conformation is crucial for RNF168-mediated H2A ubiquitination (**Extended Data Fig. 8a-b**). We anticipate that the application of activity-based chemical trapping strategies holds significant promise for the determination of other dynamic and intractable protein complexes.

Based on these results, a model describing the mechanism of RNF168-mediated H2A K13/K15 ubiquitination was proposed (**Fig. 6c**): RNF168 together with UbcH5c∼Ub engages the unmodified nucleosome, and two conformations, namely, the Ub^close^ conformation and Ub^back^ conformation were captured. In the Ub^close^ conformation, RNF168/UbcH5c∼Ub form a closed E2∼Ub conformation^39^, and our biochemical data supported that this conformation mediated productive H2A ubiquitination (**Extended Data Fig. 8b**). In the Ub^back^ conformation, the Ub motif was captured between UbcH5c and SHL DNA 3.5. The Ub^back^ conformation may provide a structural basis for the subsequent H2A K13/K15 dual-ubiquitination, and the H2A monoubiquitinated nucleosome could accommodate a second UbcH5c∼Ub without steric clashes, as evidenced by the presence of two spatially separated Ub conformations, one inside and one outside the brace of RNF168-UbcH5c (**Fig. 3e**). Supportively, disrupting the potential interfaces between Ub and UbcH5c or nucleosomal SHL DNA 3.5 by introducing mutations in Ub or UbcH5c appears to slightly impair the subsequent ubiquitination activity on the H2AUb NCP by RNF168 (see details in **Extended Data Fig. 13**).

It should be noted that structural determination of the ternary RNF168/UbcH5c-Ub/NCP complex was grueling. Direct incubation or E3-E2 linear fusion strategies all failed due to the weak, transient, and dynamic ubiquitination process of RNF168/UbcH5c on nucleosomes (**Extended Data Fig. 2**). The stable RNF168/UbcH5c/NCP complex structures we determined were based on chemical trapping strategies. We acknowledge that a previous data-driven structural modelling of RNF168/UbcH5c/nucleosome complex, based on biochemical and biophysical experiments including NMR interaction mapping, enzymatic mutagenesis and cross-linking mass spectrometry^21^, aligns with our cryo-EM determination. Our work makes advancements in several key aspects: first, we provide detailed interaction information at the residue-pair resolution for RNF168, UbcH5c, and nucleosomes. Second, our work elucidates and proposes a novel “helix-anchoring” binding mode for the monomeric E3 ligase RNF168 on the nucleosome, in contrast to the dimeric nucleosomal E3 ligases. This finding expands our knowledge of the structural and mechanistic diversity of E3 ligase-nucleosome interactions. Furthermore, we conducted biochemical investigations into the efficiency of Ub installation and the interplay between the initial and subsequent Ub modifications on the adjacent H2A K13 and K15 sites. Importantly, we structurally captured the RNF168/UbcH5c-Ub/monoubiquitinated nucleosome complex in transient H2AK13/15 adjacent dual-monoubiquitination, which provides structural evidence for that the pre-modified Ub motif on H2A does not act as a steric hindrance to the second E2∼Ub molecule and it also does not interfere with the formation of the E2∼Ub close conformation in subsequent ubiquitination event. These explorations provide a valuable framework for understanding the histone ubiquitination process of adjacent dual-monoubiquitination.

Our biochemical results, along with findings from other groups^6,7^, provide evidence suggesting that RNF168 primarily facilitates the dual-monoubiquitination of adjacent H2A K13 and K15 sites *in vitro*. However, *in vivo* studies have revealed the presence of K27-or K63-linked polyubiquitination on the H2A K13/K15 sites^17,34^, indicating the involvement of additional factors that stimulate the polyubiquitination process *in vivo*. The Ub E3 ligase RNF8 and the E2 enzyme UBC13/MMS2, which are also involved in DNA damage response signalling, have been reported to function upstream of RNF168 *in vivo*^17^. We examined the *in vitro* nucleosomal H2A ubiquitination activity of RNF168 in the presence of RNF8 and/or UBC13/MMS2, which implies a potential uncoupling between the initial monoubiquitination of H2A K13/K15 and the subsequent polyubiquitination elongation events (see more details in **Extended Data Fig. 14a-b**). Consistently, a “two-step reaction” model that two distinct E2/E3 pairs participate in the H2AK13/15 ubiquitination process has been previously proposed, in which RNF168 and UbcH5c mediate the initial monoubiquitination step that primes H2A, while RNF8 and Ubc13/Mms2 catalyze the subsequent elongation of polyubiquitin chains on the monoubiquitinated protein^6^.

Finally, our work represents the first biochemical and structural case study of adjacent dual-monoubiquitination mechanism. There is still much to investigate regarding the neighboring histone ubiquitination, such as H2A K118/119 ubiquitination by Ring1B/Bmi1^57,58^, H2A K125/127/129 ubiquitination by BRCA1/BARD1^56^ and H3 K18/23 ubiquitination by UHRF1^59^. An important example of the functional implication of the neighboring histone ubiquitination system is that dual-monoubiquitination of H3 K18 and K23 recruits and activates the maintenance DNA methyltransferase DNMT1 by interacting with its RFTS domain^60,61^. Whether there are specific readers for the H2AK13Ub/K15Ub dual-monoubiquitination remains to be investigated in further studies, which may open up new possibilities for the identification of such “bidentate” Ub sites. Future research remains to be continued on the writing, reading, and erasing “codes” of multiple mono-ubiquitinated histones or nucleosomes.

## Methods

### Cloning and plasmid construction

The cDNA of human RNF168, UbcH5c, RNF8, and SET8 were synthesized by GenScript Biotech (Nanjing, China) with sequence optimized for *Escherichia coli* overexpression. RNF168^FL^ was cloned into the pGEX6P2 vector with an N-terminal GST–HRV3C-tag, and RNF168^RING^, SET8 were individually cloned into the pET28a vector with an His_6_–HRV3C-tag. RNF8 was cloned into the pET28a vector with an N-terminal His_6_–SUMO-tag. The human Uba1, Ub (including Ub K/R mutants, Ub-G76C, and Ub-MCQ (a Cys residue insertion between the first Met residue and the second Gln residue of the Ub sequence)), human core histones (including H2A, H2B, H3(C96S/C110S), and H4) were cloned into corresponding vectors as previously described^25^. The sequence of human histone H2A(K15^C^-129) was cloned into the pCold-Trigger factor (TF) vector (Takara Bio) with a SUMO coding sequence inserted to generate His_6_–TF–SUMO–H2A(K15^C^-129) sequence. For UbcH5c–RNF168^FL^fusion protein, a sequence of UbcH5c–GSGSRS–RNF168^FL^ was cloned into the pET28a vector with an His_6_–HRV3C-tag. Fusion sequences, truncations, or mutations were generated by homologous recombination or standard site-directed PCR mutagenesis methods.

For the intein-splicing strategy, the N-terminal fragment of gp41-1 intein (IntN) was genetically fused to the C-terminus of RNF168^RING^ to generate RNF168^RING^–IntN, and the C-terminal fragment of intein gp41-1 (IntC) following a His_6_–SUMO-tag was fused to the N-terminus of the UbcH5c to generate IntC–UbcH5c. Thus, RNF168^RING^ and UbcH5c would be ligated by a 6-residue linker SGYSSS after the splicing reaction.

### Protein expression and purification

RNF168^FL^, RNF168^RING^, RNF8^RING^, RNF168^RING^–IntN, and SET8 plasmids were individually transformed into BL21(DE3) *Escherichia coli* cells. Cells were supplemented with 0.2 μM ZnCl_2_ and induced with 0.5 mM isopropyl β-D-thiogalactopyranoside (IPTG) at 16°C. RNF168^FL^ was further purified by GST affinity chromatography, HRV3C protease cleavage, ion exchange (HiTrap Heparin column, GE Healthcare), and size-exclusion chromatography (Superdex 200 10/300 GL column, GE Healthcare). RNF168^RING^ and RNF168^RING^–IntN were purified by Ni affinity chromatography, HRV3C protease cleavage, anion exchange (MonoS column, GE Healthcare), and size-exclusion chromatography (Superdex 75 10/300 GL column, GE Healthcare). UbcH5c was purified by Ni affinity chromatography, HRV3C protease cleavage, and size-exclusion chromatography (Superdex 75 10/300 GL column, GE Healthcare). IntC–UbcH5c was purified similarly, except that the His_6_–SUMO-tag was removed by ULP1 protease. RNF8^FL^ or RNF8^RING^ (351–485) was purified by Ni affinity chromatography, ULP1 protease cleavage, and size-exclusion chromatography (HiLoad 16/600 Superdex 75 prep grade column, GE Healthcare). SET8 was purified in the same way as RNF168^RING^. UbcH5c–RNF168^FL^ fusion protein was purified in a similar way as RNF168^RING^ except for the anion exchange chromatography.

Uba1, Ub (including Ub K/R mutants, Ub-G76C, and Ub-MCQ), human core histones (including H2A, H2B, H3(C96S/C110S), and H4) and mutants, human histone octamer and mutants were recombinantly expressed and purified as previously described^25^. Histone H2A(K15^C^-129) was expressed in BL21(DE3) *Escherichia coli* cells as previously described^62^ and purified by Ni affinity chromatography. After the ULP1 protease cleavage, a second Ni affinity chromatography under a denaturing condition (6 M guanidine hydrochloride (Gn·HCl) added) was performed to remove the His_6_–TF– SUMO-tag. The flowthrough containing H2A(K15^C^-129) was further purified by reversed-phase high-performance liquid chromatography (RP-HPLC) and lyophilized into powder.

### Nucleosomal DNA preparation

The DNA for nucleosome reconstitution (147-bp Widom 601 DNA) was prepared as previously described^25^. The sequence of the DNA is listed below.

147-bp Widom 601 DNA:

<u>CTGGAGAATCCCGGTGCCGAGGCCGCTCAATTGGTCGTAGACAGCTCTAGCACCGCTTAAACGCACGTACGCGCTGTCCCCCGCGTTTTAACCGCCAAGGGGATTACTCCCTAGTCTCCAGGCACGTGTCAGATATATACATCCTGT</u>

### Chemical synthesis of ubiquitinated H2A

#### Preparation of fluorescent ubiquitin-conjugated H2A variants

Three different ubiquitinated H2A variants, including histone H2A K13/K15Ub, H2A K13R/K15Ub, and H2A K13Ub/K15, were synthesized using the method previously reported by our team^24^. The chemical synthesis route is illustrated in **Supplementary Figure 1a**. Briefly, the histone H2A variant (including H2A K13/K15C, H2A K13R/K15C, H2A K13C/K15) was reacted with 2-((2-chloroethyl)amino)ethane-1-(S-acetaminomethyl)thiol, then the Acm group was removed by PdCl_2_, followed by one-pot ligation with fluorescence-labelled (Fluo)-Ub-MesNa (prepared as described below). The intermediates and final ubiquitinated H2A variants were purified by RP-HPLC, and characterized by liquid chromatography-electrospray ionization-mass spectrometry (LC-ESI-MS, **Supplementary Figure 2**).

Fluo-Ub-MesNa was prepared using the E1-mediated method^63^. In brief, Ub-MCQ was reacted with 1.2 equivalent fluorescein-5-maleimide, and the resulting Fluo-Ub was purified using a Superdex 75 size-exclusion column 10/300 GL (GE Healthcare) in buffer (20 mM HEPES, 150 mM NaCl and 1 mM DTT, pH = 7.5). Fluo-Ub was mixed in a buffer containing 2 mM ATP, 1 mM MgCl_2_, and 2 mM TCEP. Subsequently, 100 mM MesNa was added to the buffer. After adjusting the pH to 7.3, the thioesterification reaction was initiated by adding 1 µM hUBA1. The reaction proceeded at 37°C for 3 hours. The resulting product, Fluo-Ub-MesNa, was analyzed and purified using RP-HPLC.

#### Preparation of wild-type (Ub^WT^)or mutated (Ub^Mut^) ubiquitin-conjugated H2A variants

Ub(1–75)-MesNa (Ub^WT^), Ub^K6D/R42E/K48D/R72E^-MesNa, Ub^E34R/G35K^-MesNa, and Ub^D39K/R74E^-MesNa were prepared as previously described^24^. Then, each Ub-MesNa was reacted with H2A K13C-CAET-SH/K15 to generate the product H2A K13Ub^WT^/K15 or H2A K13Ub^Mut^/K15 via native chemical ligation (**Supplementary Figure 1b**). Each product was analyzed and purified using RP-HPLC, and characterized by LC-ESI-MS (**Supplementary Figure 2**). The lyophilized product was reconstituted into histone octamer and the histone octamers were characterized by SDS-PAGE (**Supplementary Figure 1c**).

### Generation of the E2–Ub–Nucleosome intermediate mimics for the intein-based E2–Ub conjugated nucleosome strategy

As shown in **Fig.2 and Extended Fig. 3**, First, we obtained H2A K15C-CAET-Ub and 5,5’-dithio bis-(2-nitrobenzoic acid) (DTNB)-activated IntC-UbcH5c (C85 only) by protein chemical synthesis, respectively, followed by a disulfide bond exchange reaction to obtain the H2A K15-CAET-Ub-IntC-UbcH5c. Then H2A K15-CAET-Ub-IntC-UbcH5c is assembled into nucleosomes.

#### The details of the process are as follows

##### Preparation of H2A K15C-CAET-Ub

Lyophilized H2A K15C peptide was dissolved in reaction buffer (100 mM HEPES, 6 M Gn ·HCl, pH = 8.5) at a final concentration of 1 mM. Then, a nucleophilic substitution reaction was initiated by the addition of 40 equiv. of CAET molecule for 4-8h at 37°C. The reaction was analysed by High-performance liquid chromatography (RP-HPLC) during the reaction and purified by RP-HPLC when the reaction was completed. Next, lyophilized H2A K15-CAET-acetaminomethyl (Acm) was dissolved in ligation buffer (100 mM NaH_2_PO_4_, 6M Gn ·HCl, pH = 7.5) at a final concentration of 1 mM. After that, 5 mg/mL of tris(2-carboxyethyl) phosphine (TCEP) and 30-fold equiv. of PdCl_2_ was added, and the pH was adjusted to 7. The deprotection reaction was carried out at 27°C for 2h to remove the Acm group. After the complete removal of the Acm group was monitored by RP-FPLC, 500 equiv. of 4-mercaptophenyl acetic acid (MPAA) was added and dissolved by sonication. Next, 1.5 equiv. of Ub(1–75)-MesNa was added, the pH was adjusted to 6.3, and the reaction was carried out at 37°C overnight. The final product of H2A K15C-CAET-Ub was purified by RP-HPLC and lyophilized.

##### Preparation of DTNB activated IntC-UbcH5c (C85 only)

Recombinantly expressed His-SUMO-IntC–UbcH5c (C85 only) was purified by Ni affinity chromatography, Ulp1 protease cleavage, and purified by RP-HPLC. The lyophilized IntC-UbcH5c (C85 only) was dissolved in activation buffer (100 mM NaH_2_PO_4_, 6M Gn·HCl, pH = 7.5) at a final concentration of 10 mg/mL. Afterward, DTNB (3 equiv.) pre-dissolved in activation buffer was added and the reaction was carried out at room temperature for 30 min, analyzed, and purified by RP-HPLC.

##### Preparation of H2A K15-CAET-Ub-IntC-UbcH5c through a disulfide exchange reaction

Lyophilized IntC-UbcH5c-TNB and H2A K15C-CAET-Ub were dissolved in disulfide ligation buffer at a final concentration of 10 mg/mL. After that, they were mixed in a 1:1 equiv. ratio and reacted for 10 min at 37°C and purified directly by RP-HPLC and lyophilized.

##### Reconstitution of H2A K15-CAET-Ub-IntC-UbcH5c octamer and Nucleosome

*H2A K15-CAET-Ub-IntC-UbcH5c* octamers and nucleosomes were assembled as previously described.

### Generation of the E3-E2–Ub–Nucleosome intermediate mimics for activity-based chemical trapping strategy

As shown in **Fig.3, Extended Fig. 6 and Extended Fig. 9**, First, we obtained IntC-UbcH5c-CAET-Ub by similar steps as described above, then IntC-UbcH5c-CAET-Ub was refolding and purified by size-exclusion chromatography (Superdex 75 10/300 GL column, GE Healthcare). The refolded IntC-UbcH5c-CAET-Ub-SH was mixed with IntN-RNF168 to initiate the protein trans-splicing reaction and finally obtained the RNF168-UbcH5c-CAET-Ub.

In parallel, we synthesized 2,2′-dipyridyl disulfide (AT_2_) -activated H2A K15C and H2A K13Ub-K15C, which were added to H2B, H3, and H4 to assemble into octamers and then added to Widom 601 DNA to assemble into nucleosomes.

#### The specific steps were as follows

##### Preparation of IntC-UbcH5c-CAET-Ub

As described above, lyophilized IntC-UbcH5c (C85 only) was dissolved in reaction buffer (100 mM HEPES, 6M Gn·HCl, pH = 8.5) at a final concentration of 15 mg/mL. The subsequent steps were similar to the preparation of H2A K15C-CAET-Ub.

##### Preparation of AT2-activated H2A K15^C^

Lyophilized H2A K15C dissolved in activation buffer (100 mM NaH_2_PO_4_, 6M Gn·HCl, pH = 7.5) at a final concentration of 0.2 mM. Afterward, 2,2′-dipyridyl disulfide (AT_2_) (40 equiv.) pre-dissolved in N, N-Dimethylformamide (DMF) was added and the reaction was carried out at room temperature for 2-4 h, analyzed, and purified by RP-HPLC.

##### Preparation of H2A(1-13^Ub^-15^C^-129)

H2A (1-K13^aG^-14)-NHNH_2_ was synthesized by microwave-assisted Fmoc-based solid-phase peptide synthesis (SPPS).

Lyophilized H2A(1-K13^aG^-14)-NHNH_2_ dissolved in reaction buffer (100 mM NaH_2_PO_4_, 6M Gn·HCl, pH=6.5). Next, 1.5 equiv. of Ub(1–75)-MesNa was added, the pH was adjusted to 6.3, and the reaction was carried out at 37°C overnight. The final product of H2A(1-K13^Ub-aG^-14)-NHNH_2_ was purified by RP-HPLC and lyophilized. The auxiliary group of H2A(1-K13^Ub-aG^-14)-NHNH_2_ was detached by trifluoroacetic acid (TFA) cocktail (TFA:H_2_O:TIPS:Phenol = 87.5:5:2.5:5, vol/vol) to afford H2A(1-K13^Ub^-14)-NHNH_2._

Lyophilized H2A(1-K13^Ub^-14)-NHNH_2_ dissolved in ligation buffer (100 mM NaH_2_PO_4_, 6M Gn·HCl, pH=2.3) and precooled to –15°C. Then, 10 equiv. NaNO_2_ pre-dissolved in the connection buffer was added to oxidize the hydrazide to acyl azide and the reaction was carried out at –15°C for 20 min. 40 equiv. MPAA was added and the pH was adjusted to 5.0. Afterward, 1.2 equiv. of H2A(K15^C^-129) dissolved in the reaction buffer (100 mM NaH_2_PO_4_, 6M Gn·HCl, pH = 2.3) were added. The pH was adjusted to 6.3 and the reaction was carried out at 27 °C overnight. The product H2A(1-13^Ub^-15^C^-129) was purified by RP-HPLC and lyophilized.

##### Preparation of DTNB-activated H2A(1-13^Ub^-15^C^-129)

The reaction steps were similar to the preparation of AT2-activated H2A K15C.

##### Refolding of IntC-UbcH5c-CAET-Ub

Lyophilized IntC-UbcH5c-CAET-Ub was dissolved unfolding buffer (20 mM HEPES, 8M Urea, 10% Glycerol, pH = 7.5) at a final concentration of 1 mg/mL. Dissolved IntC-UbcH5c-CAET-Ub was transferred to a dialyzer (D-Tube Dialyzer Maxi, MWCO 6– 8 kDa, Merck) in 300 mL of unfolding buffer. The refolding buffer (20 mM HEPES, 10% Glycerol pH = 7.5) was then slowly pumped through a peristaltic pump at a flow rate of 0.6∼1 mL/min. After the urea concentration was reduced to 1 M, it was purified by size-exclusion chromatography (Superdex 75 10/300 GL column, GE Healthcare). The peak fraction was collected and flash-frozen in liquid nitrogen

##### Preparation of RNF168-UbcH5c-Ub

IntC-UbcH5c-CAET-Ub and RNF168-IntN were mixed in a 1:2 equivalence ratio to initiate the protein splicing reaction, after 4h at 30°C. The reaction was incubated at 30 °C and aliquots were taken at t = 0, 1, 2, 3, 4h and analysed by SDS-PAGE to monitor product generation (Fig 3d). The splicing product was concentrated to a volume of ∼500 µL and loaded onto a Superose 75 10/300 GL size-exclusion column (GE Healthcare) which is pre-equilibrated with complex buffer (20mM HEPES,150mM NaCl, No DTT, 5% Glycerol). Peak fractions were analysed by SDS-PAGE and the fraction of high-purity RNF168-UbcH5c-Ub was collected, concentrated and frozen in liquid nitrogen.

### Reconstitution of histone octamers and nucleosomes

Various histone octamers and corresponding nucleosomes were reconstituted by previously published protocol^25^. For clarity, nucleosomes used in structural study, *in vitro* ubiquitination assays, and fluorescence polarization binding assays are listed in **Supplementary Table 3**.

### Fluorescence polarization binding assays

RNF168^RING^ and its mutants were diluted in a concentration range from 0.003 μM to 25 μM and mixed with 10 nM Oregon Green 488 (OG488) maleimide (Thermo Fisher Scientific)-H2B (D51C)-labeled nucleosome. Fluorescence polarization signals were measured, and the fraction bound value was calculated as previously described^25^. OriginPro 9 (OriginLab) software was used to process data, fit curves, and measure *K*_d_ value. Binding curves were visualized using GraphPad Prism 10 software.

### *In vitro* RNF168 ubiquitination assays

Ubiquitination assays were performed, quenched, and analyzed as previously described^25^ with some modifications. Reactions were incubated at 30°C or 37°C and aliquots were taken. Samples were resolved by SDA-PAGE and visualized by fluorescent signals, Coomassie Brilliant Blue (CBB), or Western blot.

For investigating enzymatic relationships of RNF168^RING^/UbcH5c and RNF8^RING^/UBC13/MMS2 in the substrate priming step (the 1st ubiquitination on nucleosomal H2A), 0.2 μM Uba1, 2 μM RNF168^RING^, 1 μM UbcH5c, 2 μM RNF8^RING^, 1 μM UBC13, 1 μM MMS2, 100 μM Ub, and 0.5 μM H2A K13R/K15 nucleosome containing fluorescent H2A were used. For investigating enzymatic relationships of RNF168^RING^/UbcH5c and RNF8^RING^/UBC13/MMS2 in Ub chain elongation on nucleosomal H2A, 0.2 μM Uba1, 2 μM RNF168^RING^, 1 μM UbcH5c, 2 μM RNF8^RING^, 1 μM UBC13, 1 μM MMS2, 100 μM Ub, and 0.5 μM H2A K13R/K15ub nucleosome containing fluorescent H2A, 2 μM Degron-ub containing fluorescent Ub (the degron sequence was the same as previous study^23^) were used.

For analyzing enzymatic relationships of RNF168^RING^/UbcH5c and SET8 in the substrate priming step, 0.2 μM Uba1, 4 μM RNF168^FL^ or RNF168^RING^, 2 μM UbcH5c, 6 μM SET8, 100 μM Ub, and 0.5 μM H2A K13R/K15 nucleosome containing fluorescent H2A were used. For analyzing enzymatic relationships of RNF168^RING^/UbcH5c and SET8 in Ub chain elongation on nucleosomal H2A, 0.2 μM Uba1, 4 μM RNF168^FL^ or RNF168^RING^, 2 μM UbcH5c, 6 μM SET8, 100 μM Ub, and 0.5 μM H2A K13R/K15ub nucleosome containing fluorescent H2A were used.

For the ubiquitination assay of wild-type RNF168^RING^, UbcH5c and its mutants on H2A nucleosomes, 0.2 μM Uba1, 1 μM RNF168^RING^ or its mutants, 1 μM UbcH5c, 100 μM Ub, and 0.5 μM H2A K129C-FITC labeled nucleosome were used. For the ubiquitination assay of wild-type RNF168^RING^ and its mutants on ubiquitinated nucleosomes, 0.5 μM Uba1, 2 μM RNF168^RING^ or its mutants, 2 μM UbcH5c, 100 μM Ub, and 0.1 μM H2A K13-K15Ub labeled nucleosome were used.

For the ubiquitination assay of histone H2A K13/K15 selectivity by RNF168^RING^, 0.1 μM Uba1, 1 μM RNF168^RING^, 1 μM UbcH5c, 50 μM Ub, and 0.15 μM H2A K129C-FITC nucleosomes (H2A K13-K15, H2A K13R-K15, H2A K13-K15R, or H2A K13R-K15R) were used.

For the comparative ubiquitination assay of RNF168 activity on ubiquitinated nucleosome or H2A nucleosomes, 0.1 μM Uba1, 1 μM RNF168^RING^, 1 μM UbcH5c, 50 μM Ub, and 0.15 μM H2A K129C-FITC nucleosomes (H2A K13-K15R, H2A K13K15R, H2A K13-K15Ub, H2A K13R-K15Ub, or H2A K13Ub-K15) were used.

For the ubiquitin chain elongation assay of RNF168 activity on nucleosome, 2 μM Uba1, 4 μM RNF168RING, 4 μM UbcH5c, 100 μM Ub, and 0.5 μM H2A K15only-K129C-FITC nucleosomes (H2A K15Ub-K129C-FITC) were used.

### Cryo-EM sample preparation

#### Capturing the Conformation of RNF168-Mediated Monoubiquitination of H2A K15 by an intein-based E2-Ub conjugated nucleosome strategy

For the RNF168-UbcH5c-Ub-H2A K15 Nucleosome complex, freshly assembled of IntC-UbcH5c-Ub nucleosome (3 µM, final concentration) was mixed with freshly purified RNF168-IntN (12 µM, final concentration) at a stoichiometric ratio of 1:4 to initiate the protein splicing reaction, after 4h at 30 °C. The splicing product was concentrated to a volume of ∼100 µL and loaded onto a Superose 6 5/150 GL size-exclusion column (GE Healthcare) which is pre-equilibrated with complex buffer (20 mM HEPES, 30 mM NaCl, No DTT). Peak fractions were pooled and concentrated to 3.3 µM. The RNF168-UbcH5c-Ub-H2A K15 Nucleosome complex was then added to an equal volume of complex buffer containing 0.2% glutaraldehyde, crosslinked for 10 minutes in a 30°C water bath and quenched by the addition of a final concentration of 100 mM Tris. Then the complex was concentrated to a volume of ∼100 µL and loaded onto a Superose 6 5/150 GL size-exclusion column (GE Healthcare). Peak fractions were pooled and concentrated to 2µM for cryo-EM sample preparation.

#### Capturing the Conformation of RNF168-Mediated Monoubiquitination of H2A K15 by an activity-based chemical trapping strategy

For the RNF168-UbcH5c-Ub-H2A K15 Nucleosome complex, RNF168-UbcH5c-Ub (4 µM, final concentration) was mixed with freshly assembled AT_2_-activated H2A15 nucleosome (2 µM, final concentration) in a 2:1 equivalence ratio to undergo an activity-based disulfide bond reaction. The reaction was incubated at 30 °C and aliquots were taken at t = 0.1, 15, 30, 45, 60, 120 min and analyzed by SDS-PAGE to monitor product generation (**Extended Data Figure 11c**). The RNF168-UbcH5c-Ub-H2A K15 Nucleosome complex was then added to an equal volume of complex buffer containing 0.2% glutaraldehyde, crosslinked for 10 minutes in a 30°C water bath and quenched by the addition of a final concentration of 100 mM Tris. Then the complex was concentrated to a volume of ∼100 µL and loaded onto a Superose 6 5/150 GL size-exclusion column (GE Healthcare). Peak fractions were pooled and concentrated to ∼2 µM for cryo-EM sample preparation.

#### Capturing the Conformation of RNF168-Mediated Adjacent Dual-Monoubiquitination of H2A K13 and 15 by an activity-based chemical trapping strategy

For the RNF168-UbcH5c-Ub-H2A K13Ub Nucleosome complex, RNF168-UbcH5c-Ub (4 µM, final concentration) was mixed with freshly assembled DTNB-activated H2A K13Ub-15C Nucleosome (2 µM, final concentration) in a 2:1 equivalence ratio to undergo an activity-based disulfide bond reaction. The following steps were similar to the preparation of the RNF168-UbcH5c-Ub-H2A K15 Nucleosome complex by an activity-based chemical trapping strategy.

### Cryo-EM Data collection and image processing

A total of 5,611 (RNF168/UbcH5c∼Ub/NCP dataset by intein-based E2-Ub-NCP strategy), 2,766 (RNF168/UbcH5c∼Ub/H2A K15^C-AT^ NCP dataset by activity-based chemical trapping strategy), 8,480 (RNF168/UbcH5c∼Ub/H2A K13Ub-K15^C-AT^ NCP dataset by activity-based chemical trapping strategy) cryo-EM images were collected on a 300 kV Titan Krios cryo-transmission electron microscope that furnished with a Gatan K3 direct detector and a GIF quantum energy filter at a pixel size of 1.074 Å. A total of 951 (RNF168/UbcH5c∼Ub/NCP dataset by E2-Ub-NCP strategy) micrographs were collected on a 200kV Tecnai Arctica microscope equipped with a K2 camera at a pixel size of 0.836 Å. For all the datasets, images were recorded at a dose of 50 electrons for 32 frames with a dose rate of 2.56 e^−^ per frame. RELION v3.1.0^64^ was used for processing all the datasets including motion correction, contrast-transfer function correction, particle manual picking or auto-picking, particle extraction (256^2^ pixels), 2D and 3D classification, mask generation, 3D-auto refinement, and postprocessing. The detailed workflows were shown in **Extended Data Figure 4, 7, 10, and Supplementary Figure 3**. The data statistics regarding the cryo-EM processing were summarized in **Supplementary Table 1**. The overall resolutions were calculated according to the Fourier shell correlation (FSC) 0.143 criteria. Local resolutions were estimated using the ResMap-1.1.4.

### Model building, refinement and validation

The crystal structures of RNF168 RING domain (PDB: 4GB0), UbcH5c (PDB: 5EGG), Ub (PDB: 1UBQ) and cryo-EM structure of nucleosome core particle (PDB: 7XD1) were downloaded from the Protein Data Bank and docked into the cryo-EM maps of the RNF168/UbcH5c-Ub/ (ubiquitinated) nucleosome complex determined by intein-based E2-Ub-NCP conjugation strategy or activity-based chemical trapping strategy. These models were then merged into one molecule in WinCoot-0.8.2^65^, and residue mutations in UbcH5c and H2A were conducted. Different cryo-EM maps sharping at different *B*-factors including the best local resolutions for nucleosome and RNF168-UbcH5c module were alternating used for model building. The non-density fitting atoms were manually deleted. The model was then subjected to several rounds of manual adjustment in WinCoot-0.8.2^65^ or real-space refinement in Phenix-1.19.2^66^ to give the final model in good stereochemistry. All the model-building, refinement, and validation statistics were summarized in **Supplementary Table 1**. The figures and panels were drawn in UCSF ChimeraX-1.6.1^67^.

## Acknowledgments

We thank the National Key R&D Program of China (No. 2022YFC3401500 for L. Liu, and 2023YFA0915300 for M. Pan) for financial support. This study was supported by the National Natural Science Foundation of China (22137005, 92253302, 22227810 for L. Liu, and 22277073 for M. Pan), Shanghai Rising-Star Program (22QA1404900), Shanghai Pilot Program for Basic Research - Shanghai Jiao Tong University (21TQ1400224)) and funding from China Postdoctoral Science Foundation (2022TQ0170 and 2022M720075 for H. Ai). L. Liu thanks XPLORER prize and the New Cornerstone Investigator Program. H. Ai thanks the funding from the National Facility for Translational Medicine (Shanghai). We acknowledge the Tsinghua University Branch of China National Center for Protein Sciences (Beijing) for cryo-EM data collection.

## Author contributions

H. Ai, Z. Tong, Z. Deng, C. Tian, L. Liu and M. Pan proposed the idea, designed the experiments, and analyzed the results. Z. Tong and L. Liu design and optimization of intein-based E2-Ub-nucleosome conjugation strategy, activity-based chemical trapping strategy. Z. Tong, Z. Deng, H. Ai, S. Tao, and Q. Shi cloned the plasmids, expressed the proteins (RNF168, UbcH5c, histones and mutants) and reconstituted the nucleosomes. Z. Tong, H. Ai, and Z. Deng synthesized the intein-E2-Ub conjugated H2A K15^C^, E2-Ub conjugated H2A K15^C^, H2A K15^C-AT^ probe, and H2A K13Ub-K15^C-AT^ nucleosome probes. H. Ai, Z. Deng, Z. Tong, and S. Tao prepared fluorescently labelled H2A histones. Q. Shi and Z. Deng synthesized the fluorescent labelled ubiquitinated histone H2A. Z. Deng and J. Liang synthesized the H2A K13Ub^Mut^/K15 histone. H. Ai performed the RNF168 H2A K13/K15 selectivity assays, and RNF168/UbcH5c mutant activity experiments on unmodified nucleosome substrates or acidic patch-mutated nucleosomes. Z. Tong performed the RNF168 ubiquitination experiments of ubiquitin chain elongation and Ub mutants on the H2A K15Ub NCP. Z. Deng performed the RNF8 and SET8-related RNF168 ubiquitination experiments, and the RNF168 mutant activity tests on ubiquitinated nucleosomes. Z. Tong prepared the cryo-EM samples. H. Ai, Z. Tong and Z. Deng checked the samples, and collected the cryo-EM data. H. Ai processed, determined the cryo-EM structures, and built the atomic models. Z. Deng, Z. Tong, and H. Ai collated the experimental data and prepared the figure panels and tables. H. Ai drafted the manuscript. H. Ai, Z. Deng, Z. Tong, C. Tian, L. Liu and M. Pan revised the manuscript. All authors read, discussed, and analyzed the manuscript. M. Pan, L. Liu and C. Tian supervised the project.

## Data availability

The cryo-EM maps and atomic model of RNF168-UbcH5c in complex with nucleosomes have been deposited in the Electron Microscopy Data Bank (EMDB) and Protein Data Bank (PDB) under accession codes: EMD-38099 and PDB-8X7I (RNF618/UbcH5c∼Ub/NCP complex by intein-based E2-Ub-nucleosome conjugation strategy), EMD-38102 (RNF618/UbcH5c∼Ub/NCP complex by E2-Ub-NCP conjugation strategy, 200 kV reconstruction), EMD-38100 and PDB-8X7J (RNF618/UbcH5c∼Ub/K15^C-AT^ NCP complex by activity-based chemical trapping strategy), EMD-38101 and PDB-8X7K (RNF618/UbcH5c∼Ub/H2A K13Ub-K15^C-AT^ NCP complex by activity-based chemical trapping strategy), respectively. Newly created materials from this study may be requested from the corresponding authors.

## Extended Data Figure Legends

**Extended Data Figure 1. RNF168 generates short ubiquitin chains on nucleosomal H2A K13/K15 without polyubiquitin chain linkage preference. a**, *In vitro* ubiquitination assay shows that RNF168^RING^ together with UbcH5c can generate short ubiquitin chains on H2A K13R/K15 NCP. Note that there is only one lysine (H2A K15) on H2A K13/K15 NCP that can be conjugated with ubiquitin and bands of H2AUb_3_ were observed, suggesting that at least three ubiquitin were conjugated to H2A K15. The PRC1^RING^ E3 ligase that site-specifically catalyzes nucleosomal H2A K118/K119 monoubiquitination was used as a negative control and it failed to generate significant H2A ubiquitination on H2A K13R/K15 NCP. **b**, *In vitro* ubiquitination assay to test the ubiquitin chain elongation activity of RNF168^RING^ or RNF168^FL^ on H2A K13R/K15Ub NCP. Note that in this nucleosome substrate, all the lysines on H2A were substituted by arginines so that ubiquitin can only be conjugated to the Ub motif at the H2A K15 site. It was observed that RNF168^RING^ or RNF168^FL^ generated H2AUb_3_, suggesting that no more than two Ub were conjugated to the Ub motif of the H2A K13R/K15Ub NCP and the ubiquitin chain elongation activity of RNF168^RING^ or RNF168^FL^ is weak. **c** and **d**, *In vitro* ubiquitination assays to investigate the ubiquitin chain linkage preference of RNF168^RING^ on H2A K13R/K15 NCP (**c**) H2A K13R/K15Ub NCP (**d**). It was observed that RNF168^RING^ generated di-ubiquitination at the H2A K15 site using donor Ub with single K/R mutation (K6R, K11R, K27R, K29R, K33R, K48R, and K63R) as well as the K27only ubiquitin (all the lysines were mutated arginines except for K27), and the efficiency is comparable to that of wild-type (WT) ubiquitin (**c**). Consistently, it was also observed that RNF168^RING^ further conjugated two Ub to the Ub motif of the H2A K13R/K15Ub NCP using Ub with single K/R mutation as well as the K27only Ub, and the efficiency is comparable to that of wild-type (WT) ubiquitin (**d**). Together, these results suggest that RNF168^RING^ generates short and mixed ubiquitin chains at H2A K15 sites with no linkage preference *in vitro*.

**Extended Data Figure 2. Summary of early unsuccessful attempts to capture the structure of RNF168/UbcH5c in complex with the nucleosome. a**, A schematic representation for the approach that directly incubates RNF168^FL^ and UbcH5c∼Ub with the unmodified nucleosome. No extra density on the nucleosome was observed using cryo-EM 2D classifications to analyze the sample obtained by this approach. **b**, Top: a schematic representation for the approach that incubates UbcH5c–RNF168^FL^ linear fusion protein with the unmodified nucleosome. Extra fuzzy density on NCP was observed using cryo-EM 2D classifications to analyze the sample obtained by this approach. Bottom: cryo-EM density maps contoured at different volume thresholds and the extra fuzzy density on the nucleosome disk surface is colored violet. **c**, Top: a schematic representation for the approach that incubates RNF168^RING^–UbcH5c–Ub with the unmodified nucleosome. RNF168^RING^–UbcH5c–Ub was generated by linking IntC–UbcH5–Ub obtained from E1-catalyzed E2–Ub conjugation with RNF168^RING^– IntN via intein-based protein *trans*-splicing reaction. Extra density on NCP was observed using cryo-EM 2D classifications to analyze the sample obtained by this approach. Bottom: different views of the cryo-EM density map. The extra density on the nucleosome disk surface is colored violet. Atomic model building of RNF168 or UbcH5c based on that extra density failed.

**Extended Data Figure 3. Design and generation of the H2A ubiquitination intermediate by intein-based E2–Ub–nucleosome conjugation strategy, related to Figure 2a-d. a. Top:** a detail synthetic route of the H2A K15-CAET-Ub-IntC-UbcH5c **Bottom:** intein-mediated protein trans-splicing reaction to obtain the RNF168-UbcH5c-Ub NCP complex. **b.** MS traces and deconvoluted MS spectra for purified H2A K15C-CAET-Acm, IntC-UbcH5c (C85 only), H2A K15C-CAET-SH, IntC-UbcH5c-TNB, H2A K15C-CAET-Ub, H2A K15-CAET-Ub-IntC-UbcH5c, respectively.

**Extended Data Figure 4. Cryo-EM data processing for RNF168/UbcH5c**–**Ub/NCP complex achieved by intein-based E2**–**Ub**–**nucleosome conjugation strategy. a**, A representative micrograph. **b**, CTF estimation of the micrograph shown in **a**. **c**, Representative 2D classifications. **d**, Data processing flowcharts. **e**, Euler angle distributions of the final particles for cryo-EM reconstruction. **f**, Local resolution colored cryo-EM density map. **g**, Gold standard Fourier shell correlation (FSC) curves showing the overall resolution of 3.27 Å and 3.62 Å for the final density map of RNF168/UbcH5c–Ub/NCP, and RNF168/UbcH5c, respectively.

**Extended Data Figure 5. Different linking sequences of Ub, H2A and UbcH5c, and the absence of the intein linkage between RNF168 and UbcH5c give a conserved RNF168, UbcH5c and Ub conformation on nucleosomes. a,** Synthesis starts from H2A, with intein-based linker between RNF168 and UbcH5c. **b,** Synthesis starts from UbcH5c, without intein-based linker. **c,** the cryo-EM comparison of the two maps determined in **a** and **b**, indicating a similar conformation of RNF168, UbcH5c and Ub on nucleosomes.

**Extended Data Figure 6. Design and generation of the H2A dual-monoubiquitination intermediate by activity-based chemical trapping strategy, related to Figure 3a-d. a.** (Left) Failed synthetic routes to by reaction between histone H2A(1-K13^Ub^K15^C^-129) and activated UbcH5c-CAET-Ub-TNB, no product was detected in the chromatogram of RP-HPLC analysis. **b**. Detailed synthetic route for obtaining the RNF168/UbcH5c-Ub/H2A K13Ub nucleosome complex based on an activity-based chemical trapping strategy. **c.** MS traces and deconvoluted MS spectra for purified H2A(1-K13^Ub^-K15^C^-129) and H2A(1-K13Ub-K15^C-AT^-129).

**Extended Data Figure 7. Cryo-EM data processing for RNF168/UbcH5c**–**Ub/H2A K13Ub NCP complex achieved by activity-based chemical trapping strategy. a**, A representative micrograph. **b**, Representative 2D classifications. **c**, Data processing flowcharts. **d**, Euler angle distributions of the final particles for cryo-EM reconstruction of RNF168/UbcH5c/two Ub conformation. **e**, Gold standard Fourier shell correlation (FSC) curves showing the overall resolution of 6.25 Å and 3.20 Å for the final density map of RNF168/UbcH5c/two Ub conformation, and RNF168/UbcH5c/NCP conformation, respectively. **f**, Euler angle distributions of the final particles for cryo-EM reconstruction of RNF168/UbcH5c/NCP conformation.

**Extended Data Figure 8. Analysis of the canonical closed E2∼Ub conformation and E3-E2 interfaces. a**, Alignment of four E3/E2–Ub complexes (PDB: 4AUQ, 4AP4, 6TTU, and 8GRM) that contain UbcH5 family E2s with Ub at the close conformation and the current RNF168/UbcH5c structure. The Cαs of UbcH5 E2 residues that play important roles in mediating closed E2∼Ub conformation (I88, R90, L97, L104, D116, and D117) and non-covalent ubiquitin binding stimulation (S22) are colored red and blue, respectively. **b**, *In vitro* ubiquitination assay using WT-H2A NCP to investigate the effects of UbcH5c mutants on overall nucleosomal H2A ubiquitination pattern. Data show the mean ± SD (bars) from *n* = 3 independent biological replicates. **c**, Alignment of our RNF168/UbcH5c in our complex structure with four nucleosomal E3/E2 complexes (PDB: 4R8P, 7JZV, 8IEG, and 8IEJ), highlighting the conserved E3/E2 spatial arrangement and the E3–E2 interface.

**Extended Data Figure 9. Design and generation of the H2A mono-ubiquitination intermediate by activity-based chemical trapping strategy. a.** Detailed route for obtaining the RNF168-UbcH5c-Ub NCP complex based on an activity-based chemical trapping strategy. **b.** MS traces and deconvoluted MS spectra for purified IntC-UbcH5c-CAET-Acm, IntC-UbcH5c-CAET-SH, IntC-UbcH5c-CAET-Ub-SH, H2A K15^C-AT^, respectively. **c.** A coomassie-stained SDS–PAGE gel of IntC-UbcH5c-Ub, RNF168-Int N, and aliquots of the protein splicing reaction using IntC-UbcH5c-Ub and RNF168-Int N taken at 0.01, 1, 2, 3, and 4 h. **d.** A gel filtration chromatogram of RNF168-UbcH5c-Ub obtained by protein splicing reaction. Red boxes indicate high-purity RNF168-UbcH5c-Ub used for cryo-EM sample preparation. **e.** A coomassie-stained SDS–PAGE gel of purified RNF168-UbcH5c-Ub.

**Extended Data Figure 10. Cryo-EM data processing for RNF168/UbcH5c**– **Ub/NCP complex achieved by activity-based chemical trapping strategy. a**, A representative micrograph. **b**, Representative 2D classifications. **c**, Data processing flowcharts. **d**, Cryo-EM density map at an overall resolution of 6.87 Å representing the closed E2∼Ub conformation. **e**, Local resolution colored cryo-EM density map. **f**, Gold standard Fourier shell correlation (FSC) curves showing the overall resolution of 3.39 Å and 3.27 Å for the final density map of RNF168/UbcH5c–Ub/NCP, and RNF168/UbcH5c/NCP, respectively.

**Extended Data Figure 11. Capturing the complex of RNF168-UbcH5c-mediated nucleosome monoubiquitination using activity-based chemical trapping strategy. a. Schematic representation of activity-based chemical trapping strategy.** The detailed synthetic route is shown in **Extended Data Figure 9a**. **b.** The dashed box shows the designed intermediate structure by activity-based chemical trapping strategy, which highly mimics the transition state of the substrate ubiquitination by RNF168-UbcH5c. **c.** A Coomassie-stained non-reducing SDS–PAGE gel of reconstituted H2A K15^C-AT^ nucleosome, purified RNF168-UbcH5c-Ub, and aliquots of activity-based chemical trapping reaction using RNF168-UbcH5c-Ub and H2A K15^C-AT^ nucleosome taken at 10 s,15 min, 30 min, 45 min, 60 min, 120 min. According to the molecular weight from the largest to the smallest, the bands on the gel are according to the following, respectively: RNF168-UbcH5c-Ub-H2A (53 kDa), RNF168-UbcH5c-Ub (38 kDa), H3 (15 kDa), H2A K15^C-AT^ (14 kDa), H2B (13 kDa) H4 (11 kDa). **d.** The 3.27 Å cryo-EM density map of RNF168/UbcH5c–Ub/NCP complex obtained by activity-based chemical trapping strategy. The map of RNF168/UbcH5c–Ub/NCP complex was sharpened using a B factor of −30 Å^2^ and contoured at a level of 0.011. **e.** The 3.44 Å cryo-EM density map of RNF168/UbcH5c/NCP complex. The map of RNF168/UbcH5c–Ub/NCP complex was sharpened using a B factor of −30 Å^2^ and contoured at a level of 0.011. **f**, Cryo-EM reconstruction for the E3/E2–Ub closed conformation obtained from the dataset of RNF168/UbcH5c–Ub/NCP complex achieved by activity-based chemical trapping strategy, as shown in **Extended Data Figure 10d**. **g**, Close-up view of the RNF168/UbcH5c–Ub density with models of RNF168 and UbcH5c fitted, highlighting the density of Ub^close^ located in the brace of RNF168 and UbcH5c.

**Extended Data Figure 12. Sample densities for the cryo-EM reconstructions of RNF168/UbcH5c-mediated nucleosomal H2A ubiquitination. a**–**f**, Main chain traces of the whole complex (**a**), DNA (**b**), histone octamer (**c**), RNF168 (**d**), UbcH5c (**e**), and Ub (**f**) fit into the cryo-EM density map of the RNF168/UbcH5c–Ub/NCP complex achieved by intein-based E2–Ub–nucleosome conjugation strategy. **g** and **h**, Two views of the representative region of the cryo-EM density map of the RNF168/UbcH5c–Ub/NCP complex achieved by intein-based E2–Ub–nucleosome conjugation strategy for the RNF168–acidic patch interface with side chains of RNF168 R57, S60, and R63, and some H2A/H2B residues shown. **i**–**n**, Main chain traces of the whole complex (**i**), DNA (**j**), histone octamer (**k**), RNF168 (**l**), UbcH5c (**m**), and Ub (**n**) fit into the cryo-EM density map of the RNF168/UbcH5c–Ub/NCP complex achieved by activity-based chemical trapping strategy. **o** and **p**, Two views of the representative region of the cryo-EM density map of the RNF168/UbcH5c–Ub/NCP complex achieved by activity-based chemical trapping strategy for the RNF168–acidic patch interface with side chains of RNF168 S60 and R63, and some H2A/H2B residues shown. **q**–**u**, Main chain traces of the whole complex (**q**), DNA (**r**), histone octamer (**s**), RNF168 (**t**), and UbcH5c (**u**) fit into the cryo-EM density map of the RNF168/UbcH5c–Ub/H2A K13Ub NCP complex achieved by activity-based chemical trapping strategy. **v** and **w**, Two views of the representative region of the cryo-EM density map of the RNF168/UbcH5c–Ub/H2A K13Ub NCP complex achieved by activity-based chemical trapping strategy for the RNF168–acidic patch interface with side chains of RNF168 R57, S60, and R63, and some H2A/H2B residues were shown.

**Extended Data Figure 13. Biochemical investigations of the role of the structurally observed Ub^back^ in the RNF168-mediated H2A ubiquitination. a, Models showing** the spatial proximity of SHL DNA 3.5 and Ub^back^ and the potential interface of UbcH5c and Ub^back^. The models are from the RNF168/UbcH5c–Ub/NCP complex achieved by intein-based E2–Ub–nucleosome conjugation strategy. Nucleosomal DNA is present as cryo-EM density surface and Ub^back^, UbcH5c, and H2A are shown as ribbon models. RNF168, H2B, H3, and H4 are hidden for clarity. The Cαs of Ub^back^ positive residues near the SHL DNA 3.5 (K6, R42, K48, and R72) and Ub^back^ residues near UbcH5c (E34, G35, D39, and R74) are colored red, light blue, and medium blue, respectively. **b**, Schematic representation of biochemical assays to investigate the effects of Ub^back^ mutations in the second H2A ubiquitination step. Note the Ub^back^ mutations are introduced to the H2A K13Ub/K15 NCP which is the substrate in the 2nd H2A ubiquitination step. **c**, *In vitro* ubiquitination assays described in **b** using H2A K13Ub^Mut^/K15 NCP as substrates. Coomassie Brilliant Blue-stained bands of H2AUb (substrate) and H2AUb_2_ (product) are indicated, and the band of H4 in each lane was used as a control to calculate the H2AUb_2_ generation efficiency in **d**. **d**, Quantified H2AUb_2_ generation efficiency in **c**. Combination mutations of Ub^back^ positive residues near the SHL DNA 3.5 (K6D, R42E, K48D, and R72E) and Ub^back^ residues near UbcH5c (D39K and R74E) slightly reduced the second Ub conjugation efficiency at the H2A K15 site on the H2A K13Ub^Mut^/K15 NCPs. **e**, Model showing the potential interface of UbcH5c and Ub^back^. The model is from the RNF168/UbcH5c–Ub/NCP complex achieved by intein-based E2–Ub–nucleosome conjugation strategy. The rectangle region indicates the close-up view in **f**. **f**, Close-up view showing the spatial proximity of UbcH5c and Ub^back^ and the potential interacting residues on UbcH5c and Ub^back^. **g**, Schematic representation of biochemical assays to investigate the effects of UbcH5c mutations in the first and second H2A monoubiquitination step. **h** and **i**, Left: *In vitro* ubiquitination assays described in **g** using H2A K13/K15R NCP (**h**) or H2A K13R/K15 NCP (**i**) as the substrate to test the effects of UbcH5c mutations in the first H2A monoubiquitination step. Right: Quantified H2A monoubiquitination in the left panel. Data show the mean ± SD (bars) from *n* = 3 independent biological replicates. **j**, Left: *In vitro* ubiquitination assays described in **g** using H2A K13/K15Ub NCP as the substrate to test the effects of UbcH5c mutations in the second H2A monoubiquitination step. Right: Quantified H2A ubiquitination in the left panel. Data in **d**, **h**, **i**, and **j** show the mean ± SD (bars) from *n* = 3 independent biological replicates. Note: the ubiquitinated histone H2A bearing Ub mutant were chemically synthesized as illustrated in **Supplementary Figure 1b-c and 2**.

**Extended Data Figure 14. Biochemical investigations of roles of E3 ligase RNF8 in the RNF168-mediated H2A ubiquitination. a**, *In vitro* ubiquitination assays using H2A K13R/K15 NCP as the substrate to test the nucleosomal H2A ubiquitination activity of RNF168/UbcH5c or RNF8/UBC13/MMS2. RNF168/UbcH5c efficiently ubiquitinated H2A (lane 1) while RNF8/UBC13/MMS2 failed to ubiquitinate nucleosomal H2A with or without RNF168 (lanes 2 *vs.* 3; fluorescent H2A gel image). However, the Coomassie Brilliant Blue (CBB)-stained gel image shows that RNF8/UBC13/MMS2 generated polyubiquitin chains that were not conjugated on H2A (lanes 2 *vs.* 3). **b**, *In vitro* ubiquitination assays using H2A K13R/K15Ub NCP as the substrate to test whether RNF8/UBC13/MMS2 can elongate polyubiquitin chains on the Ub motif at the H2A K15 site. UBC13/MMS2 elongated polyubiquitination chain on the H2A K13R/K15Ub NCP and RNF8 promoted this process (lanes 3 *vs.* 4, fluorescent H2A gel image). RNF168 seemed not to promote the RNF8/UBC13/MMS2-mediated polyubiquitin chain elongation (lanes 2 *vs.* 3, lanes 1 *vs.* 4; fluorescent H2A gel image), suggesting that RNF168 mediated H2A monoubiquitination and RNF8-mediated polyubiquitin chain elongation are independent. A fluorescent degron-Ub was used as a control substrate to show that RNF8/UBC13/MMS2 also elongated polyubiquitin chains on degron-Ub, suggesting that the ubiquitin chain elongating activity of RNF8 is not nucleosome context-specific (lanes 4 *vs.* 6, fluorescent H2A gel image).

