## Supplemental Figures for "Capturing Snapshots of Nucleosomal H2A K13/K15 Ubiquitination Mediated by the Monomeric E3 Ligase RNF168"

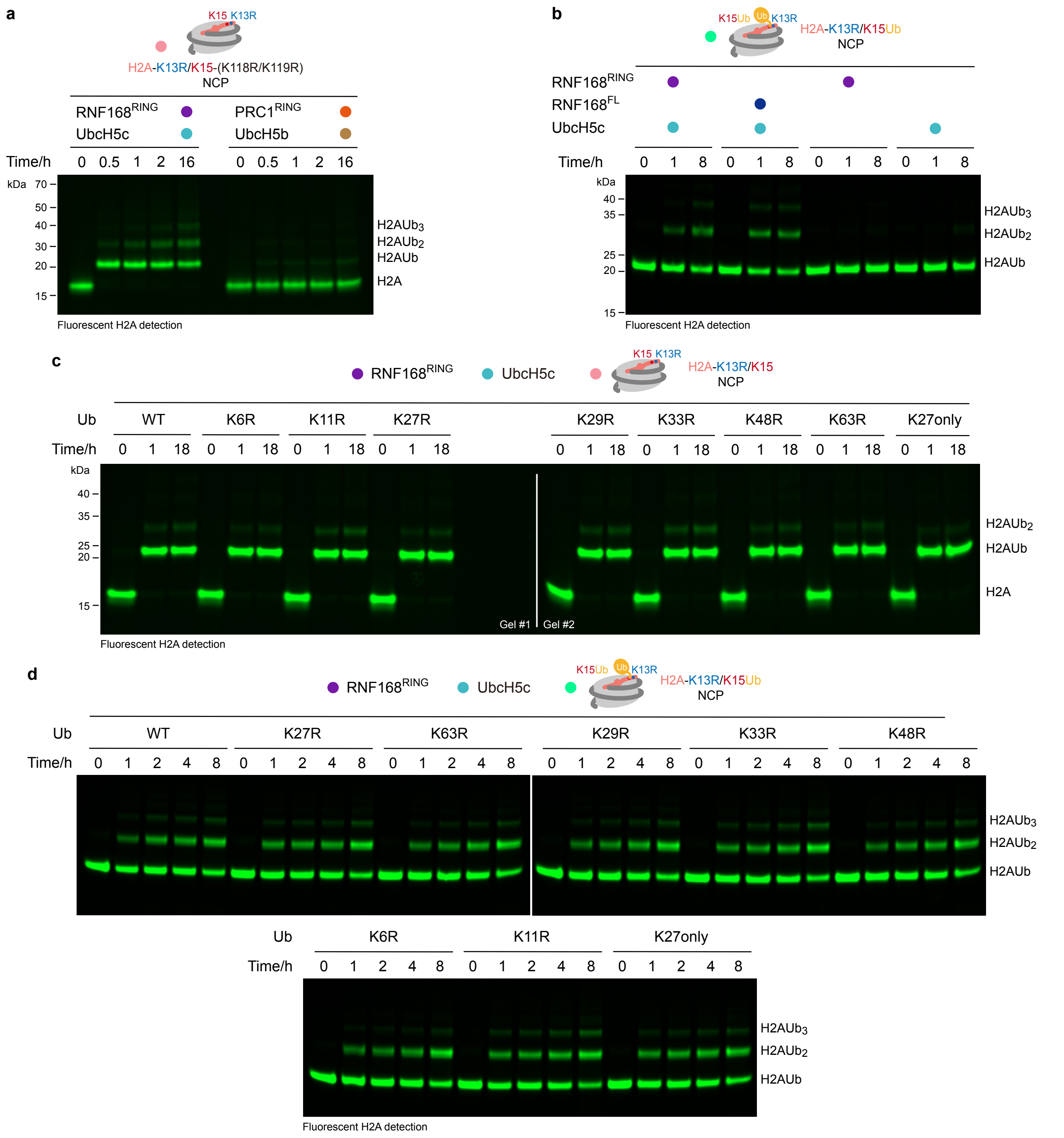


**Extended Data Figure 1. RNF168 generates short ubiquitin chains on nucleosomal H2A K13/K15 without polyubiquitin chain linkage preference.**

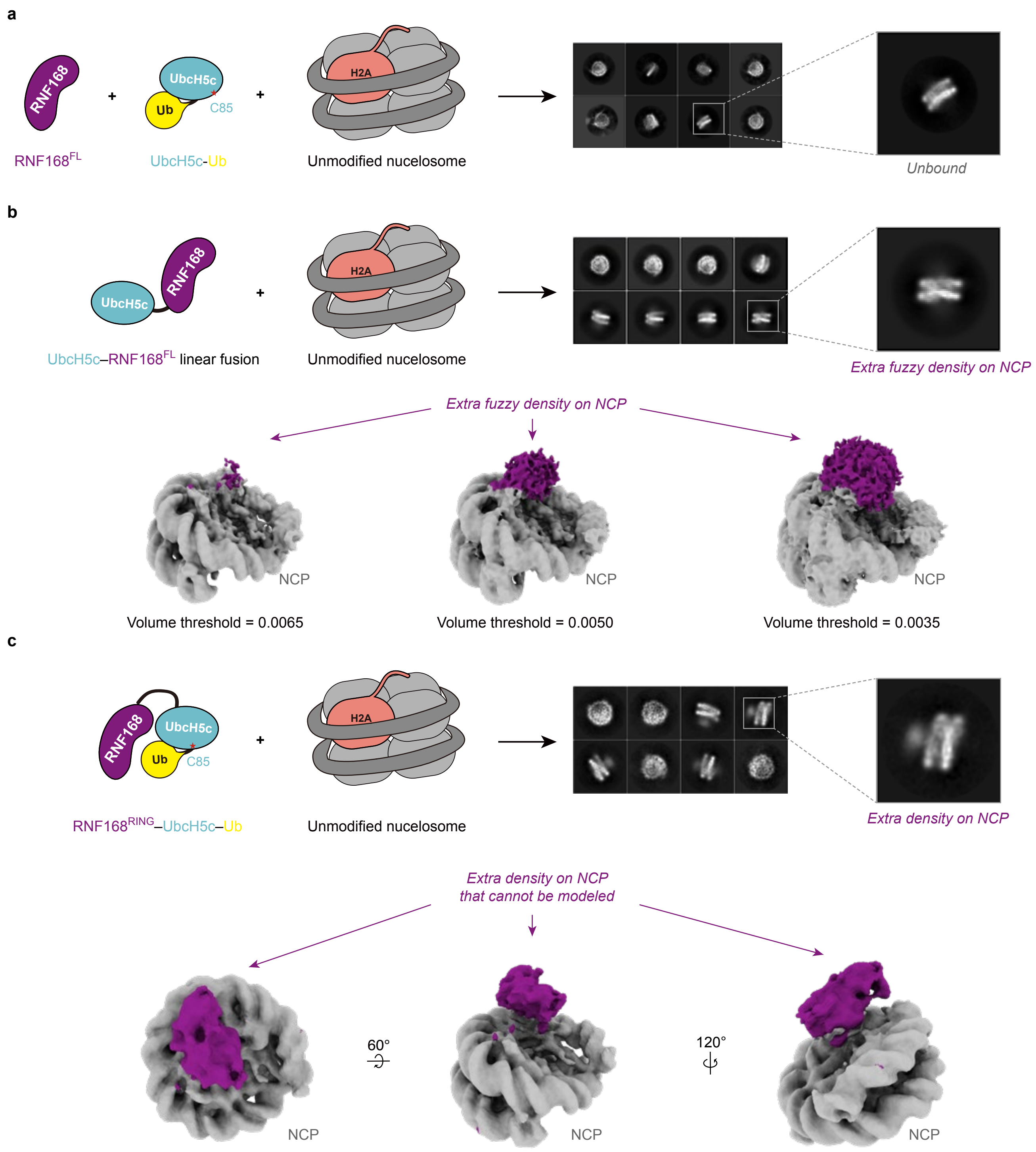


**Extended Data Figure 2. Summary of early unsuccessful attempts to capture the structure of RNF168/UbcH5c in complex with the nucleosome.**

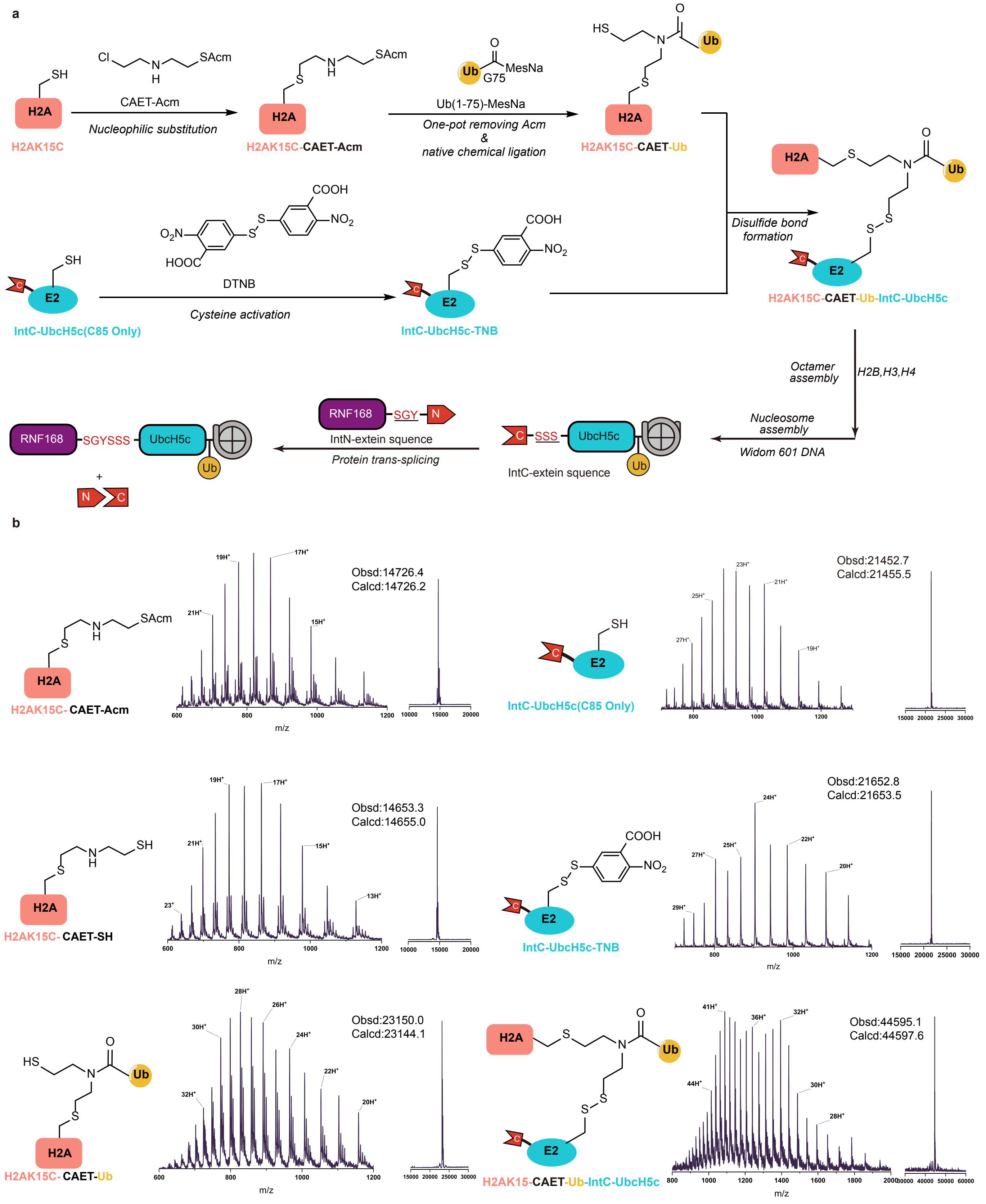


**Extended Data Figure 3. Design and generation of the H2A ubiquitylation intermediate by intein-based E2–Ub–nucleosome conjugation strategy, related to Figure 2a-d.**

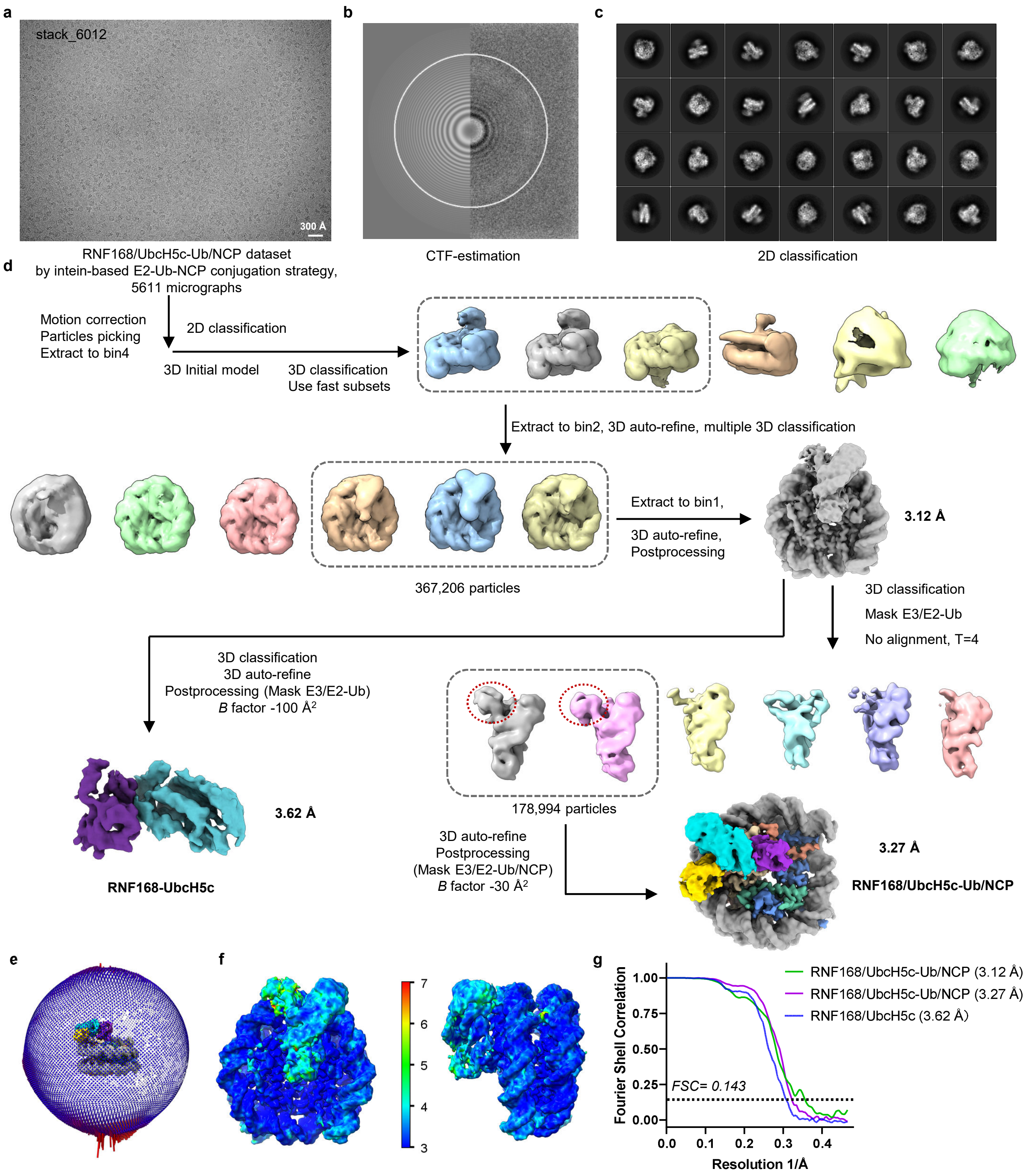


**Extended Data Figure 4 | Cryo-EM processing of RNF168/UbcH5c-Ub/NCP complex achieved by intein-based E2~Ub-nucleosome conjugation strategy**

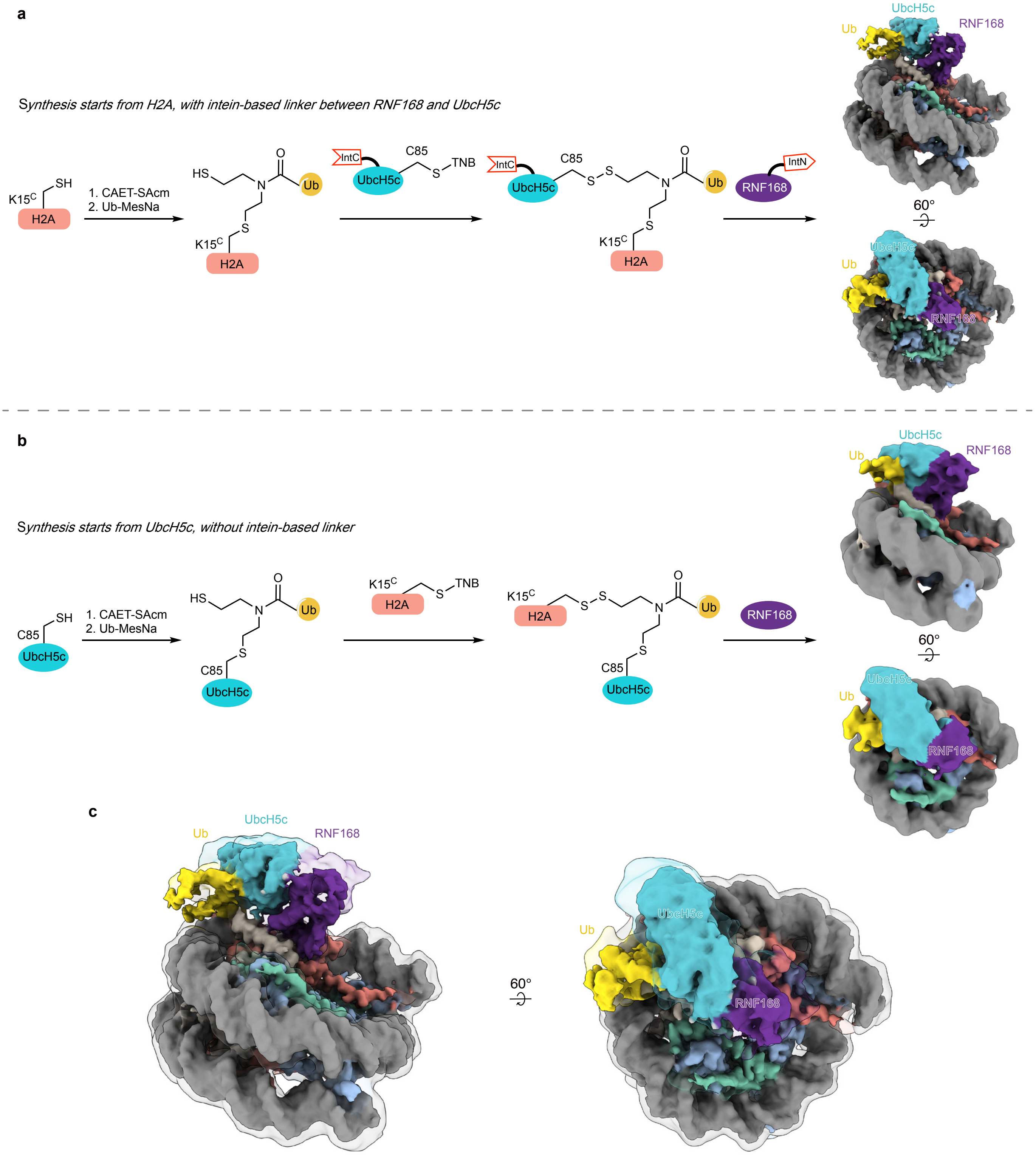


**Extended Data Figure 5. Different linking sequences of Ub, H2A and UbcH5c, and the absence of the intein linkage between RNF168 and UbcH5c give a conserved RNF168, UbcH5c and Ub conformation on nucleosomes.**

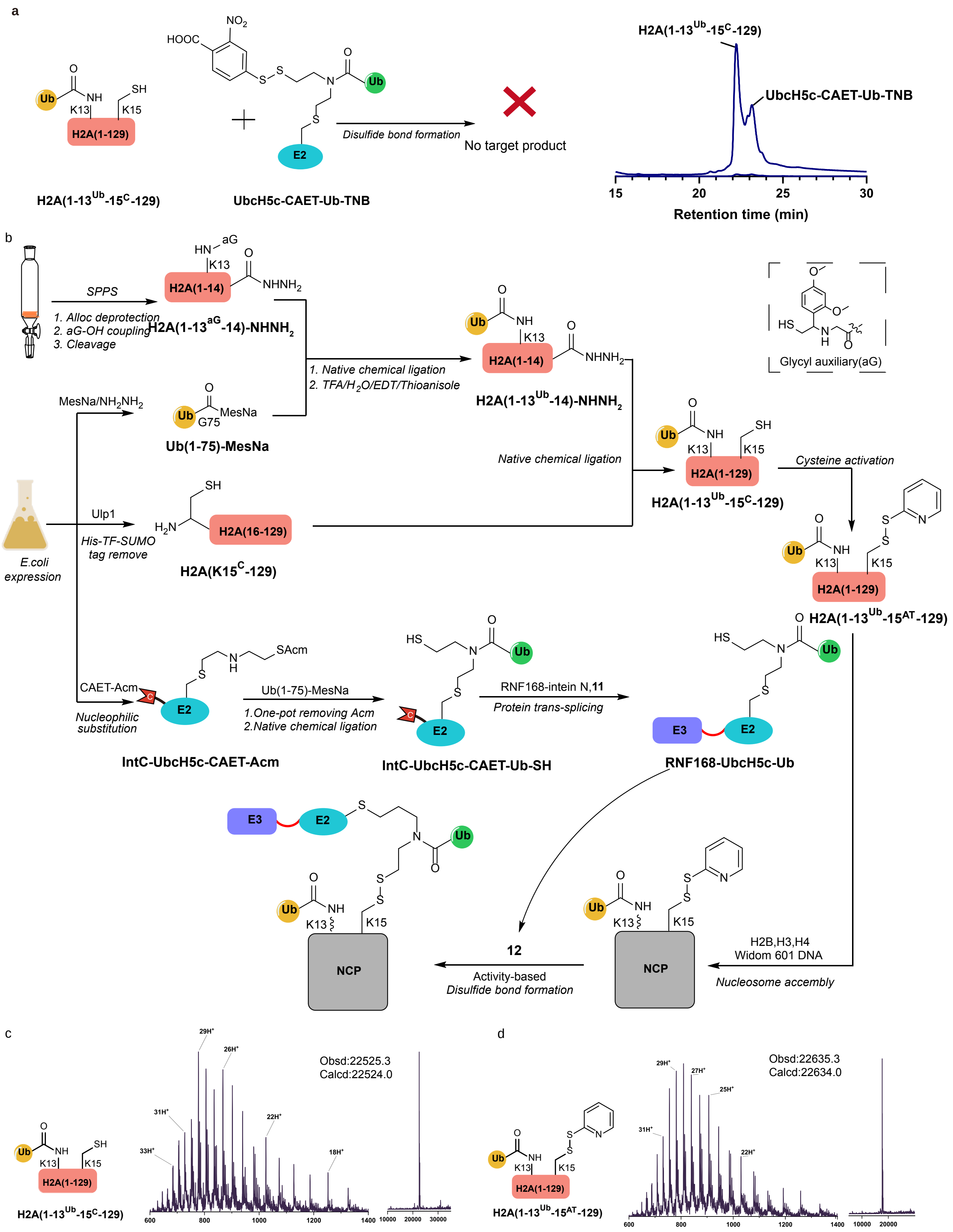


**Extended Data Figure 6. Design and generation of the H2A dual-monoubiquitylation intermediate by activity-based chemical trapping strategy, related to Figure 3a-d.**

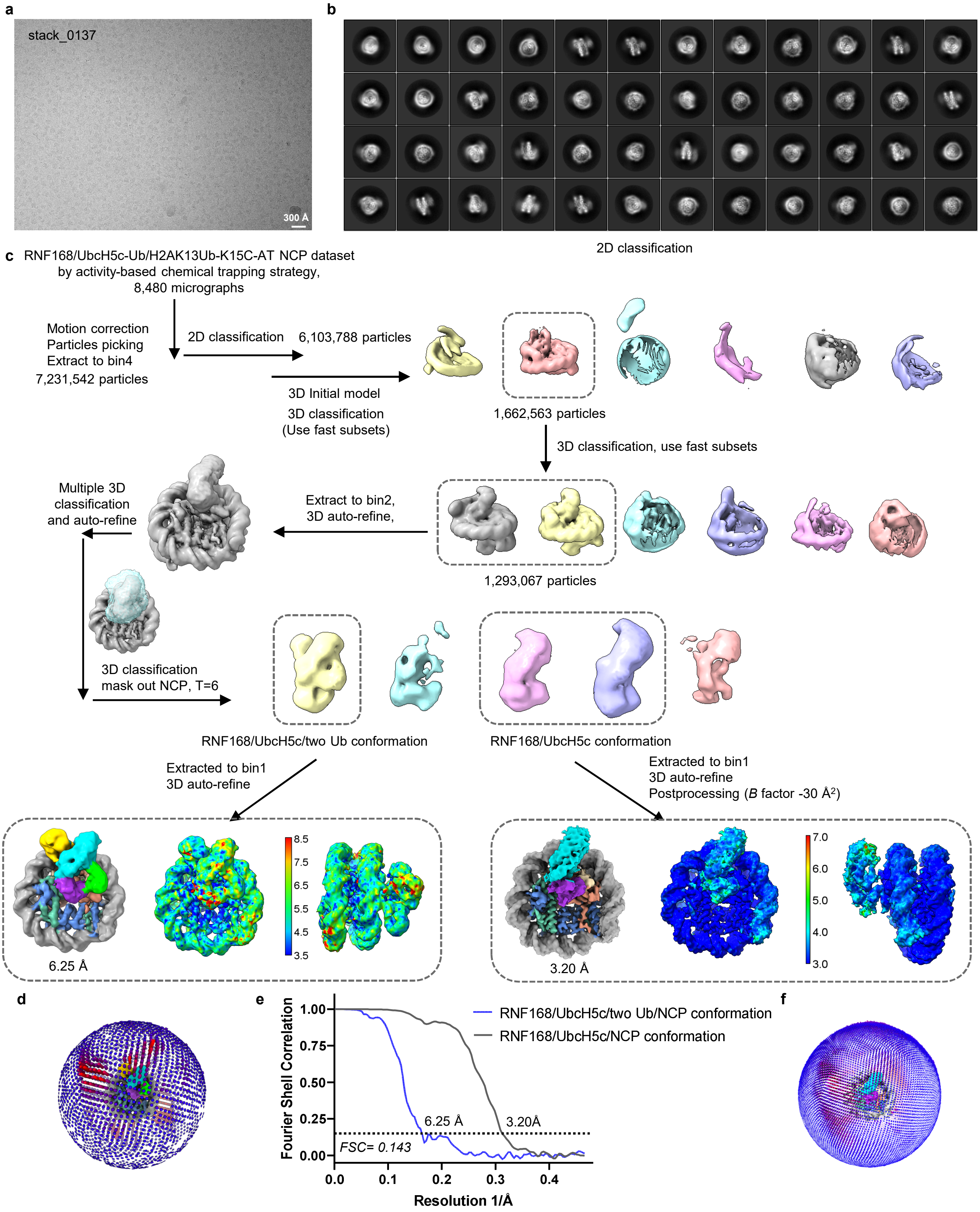


**Extended Data Figure 7. Cryo-EM data processing for RNF168/UbcH5c–Ub/H2AK13Ub NCP complex achieved by activity-based chemical trapping strategy.**

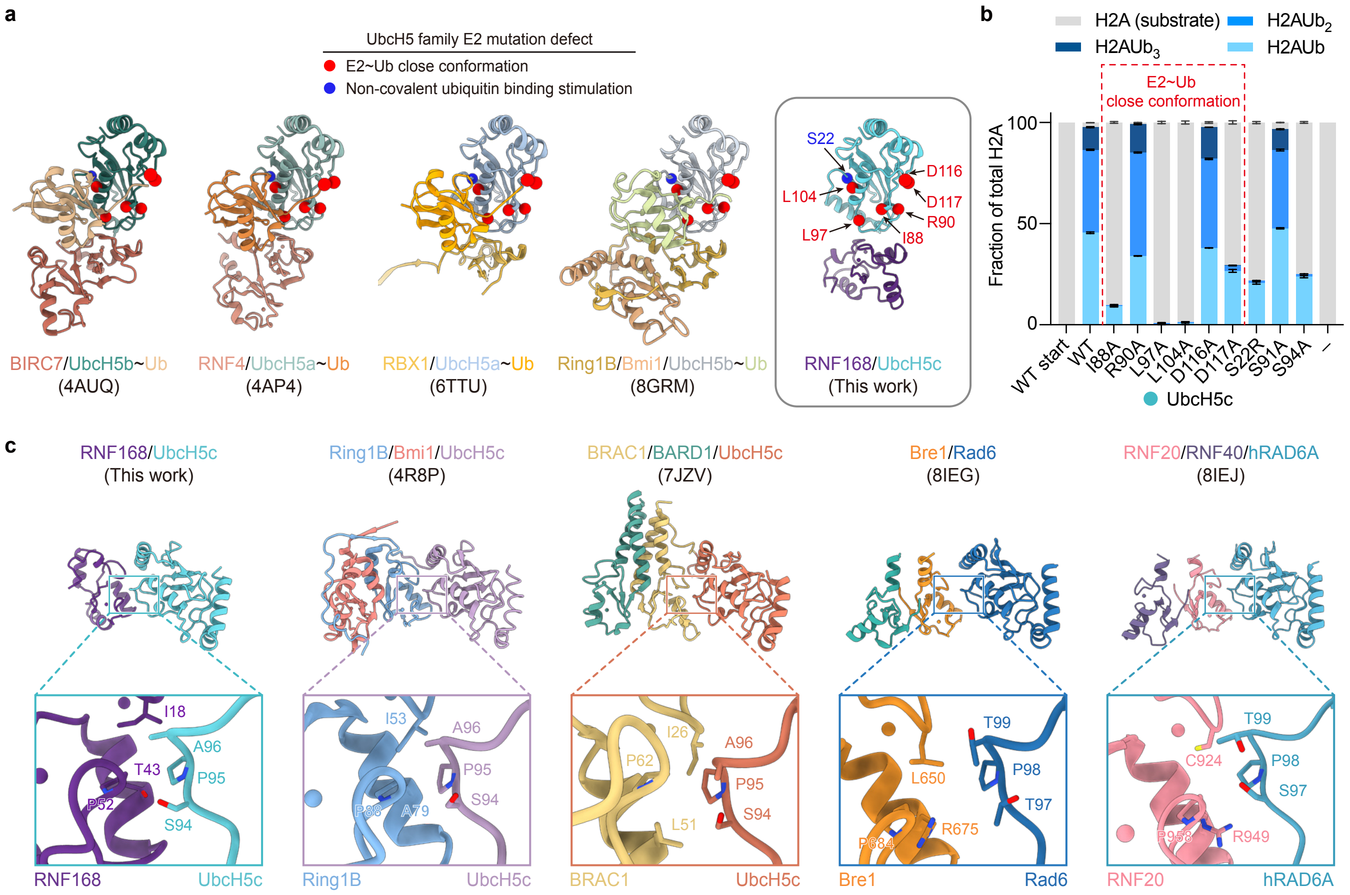


**Extended Data Figure 8. Analysis of the canonical E2~Ub close conformation and E3-E2 interfaces.**

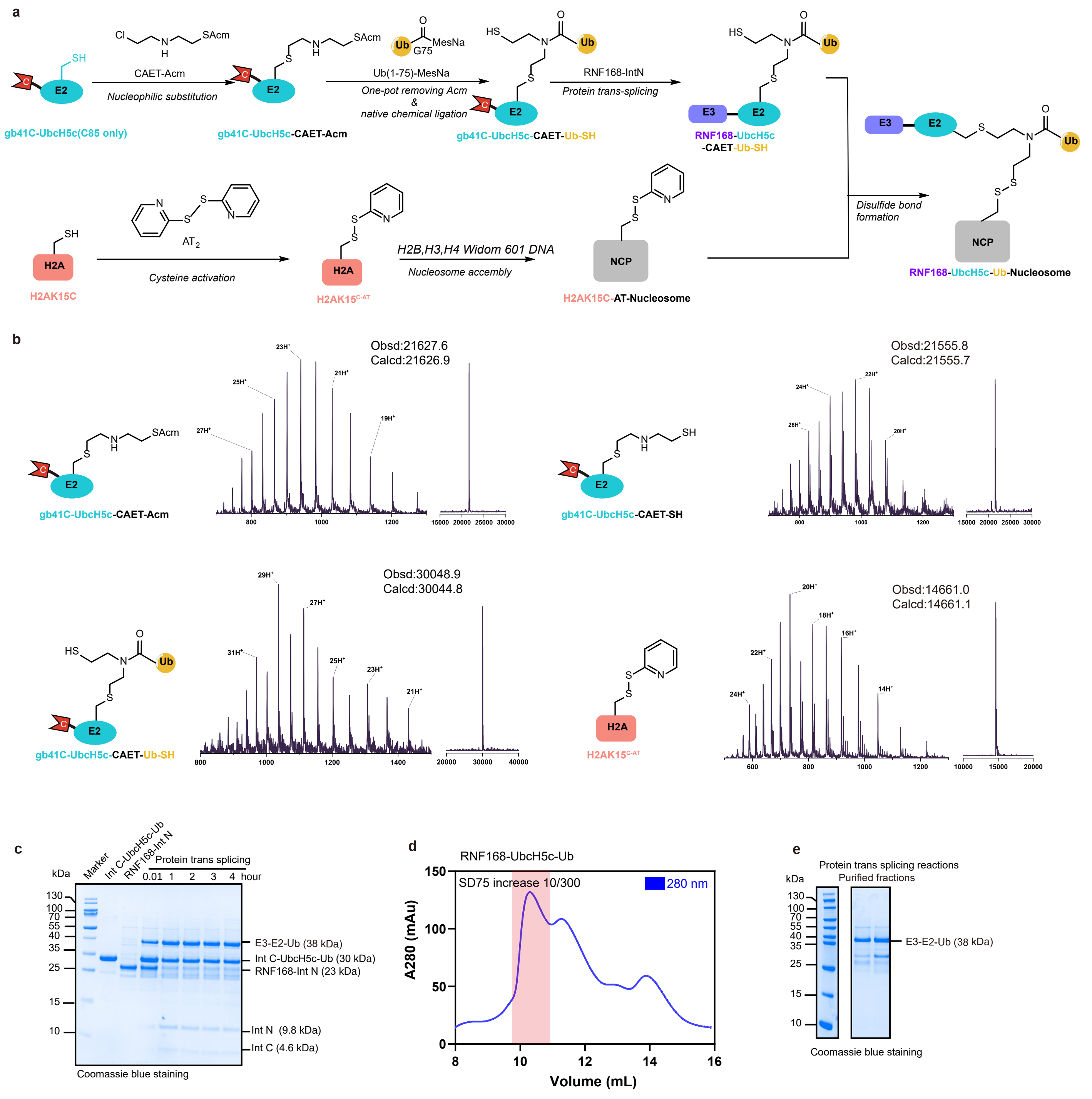


**Extended Data Figure 9. Design and generation of the H2A ubiquitylation intermediate by activity-based chemical trapping strategy.**

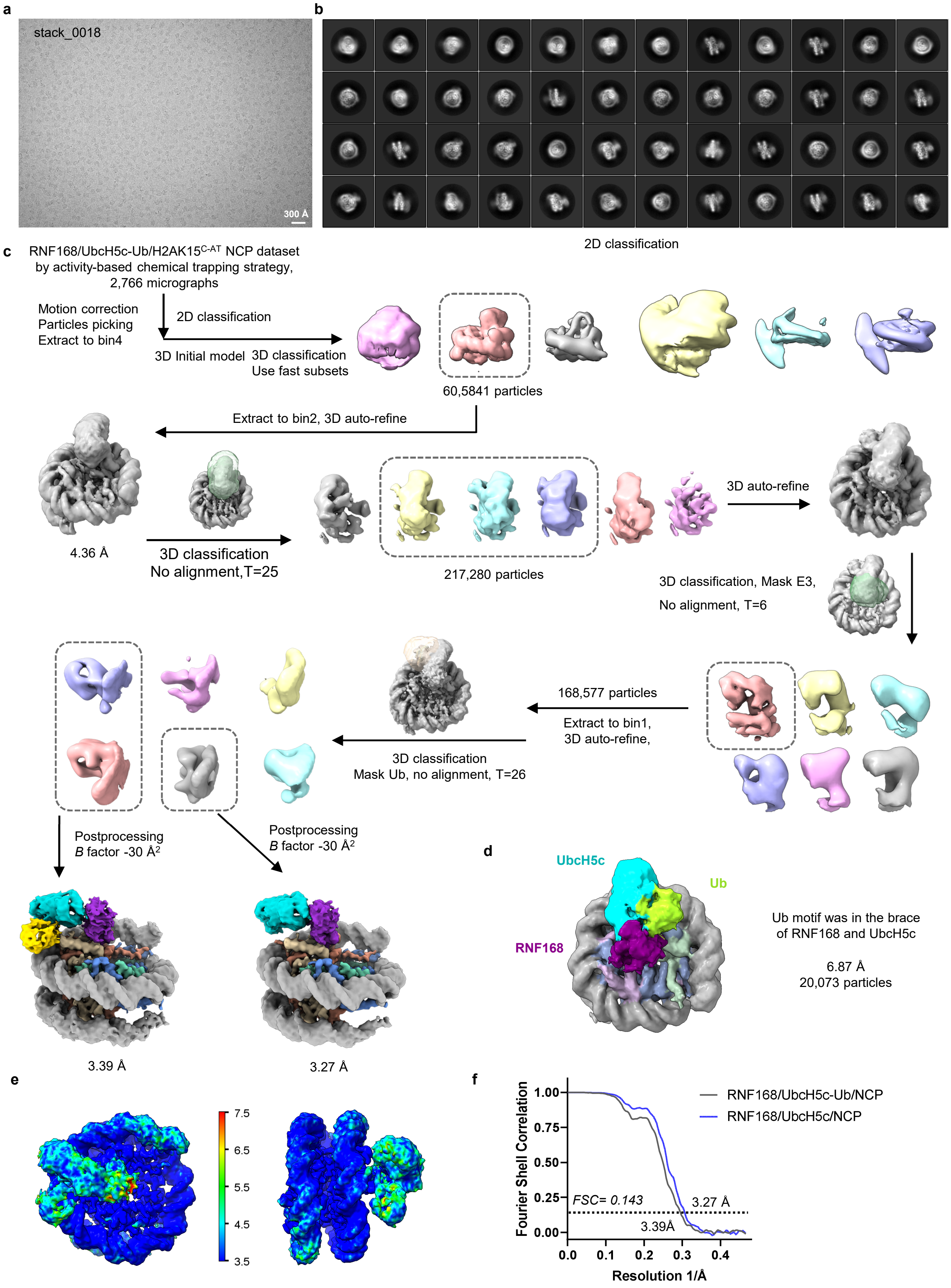


**Extended Data Figure 10. Cryo-EM data processing for RNF168/UbcH5c–Ub/NCP complex achieved by activity-based chemical trapping strategy.**

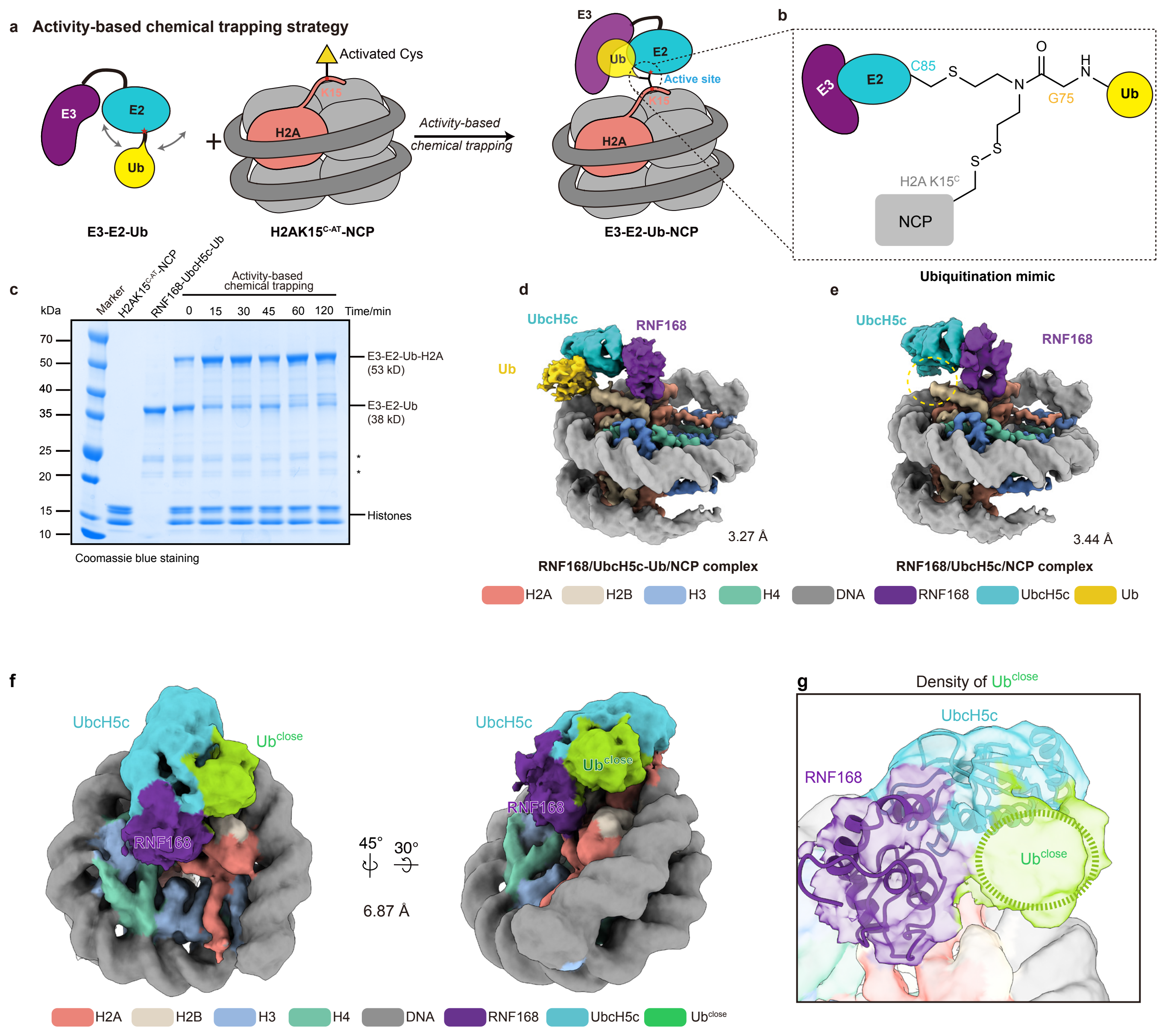


**Extended Data Figure 11. Capturing the complex of RNF168-UbcH5c-mediated nucleosome monoubiquitination using activity-based chemical trapping strategy.**

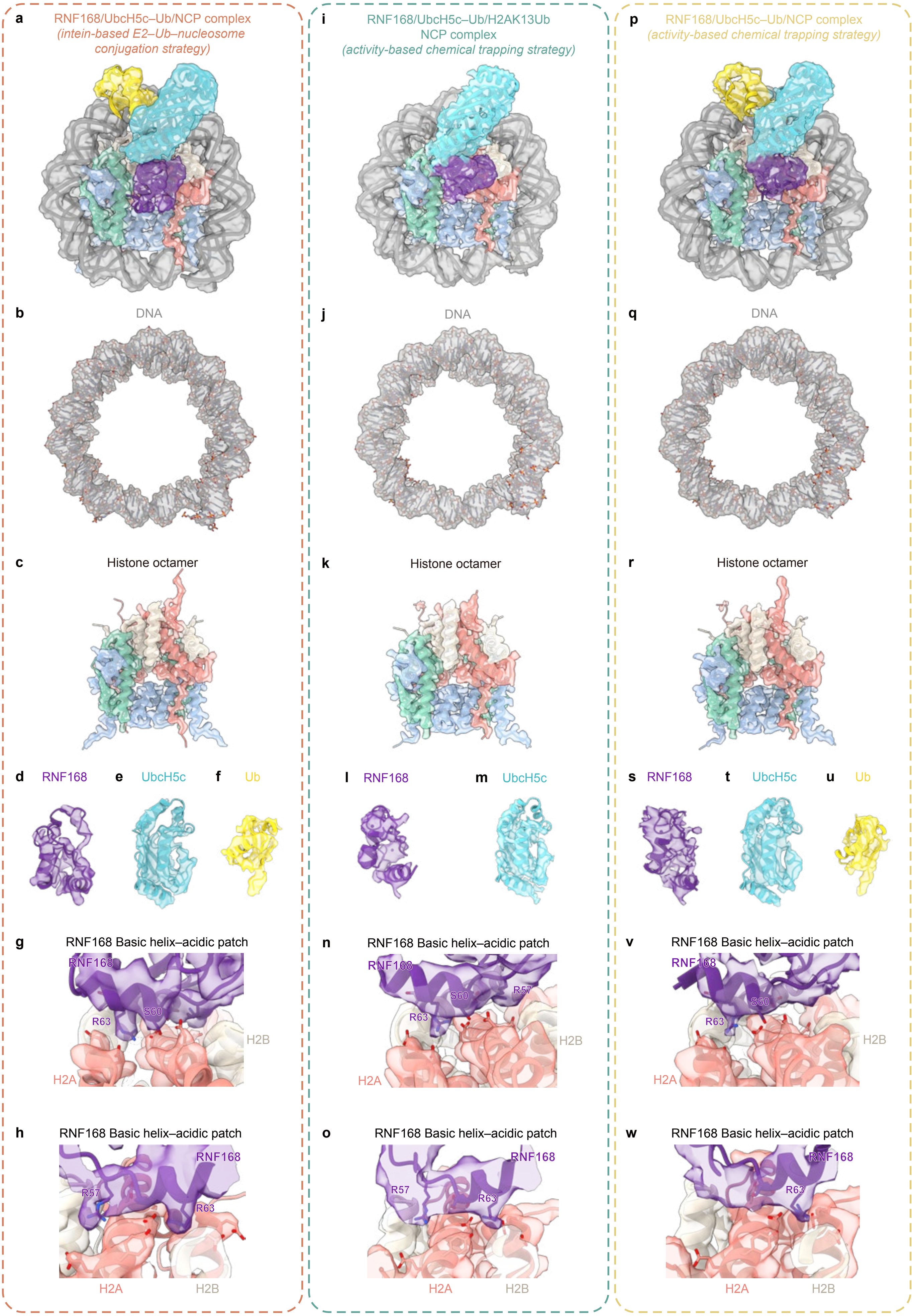


**Extended Data Figure 12. Sample densities for the cryo-EM reconstructions of RNF168/UbcH5c-mediated nucleosomal H2A ubiquitinations.**

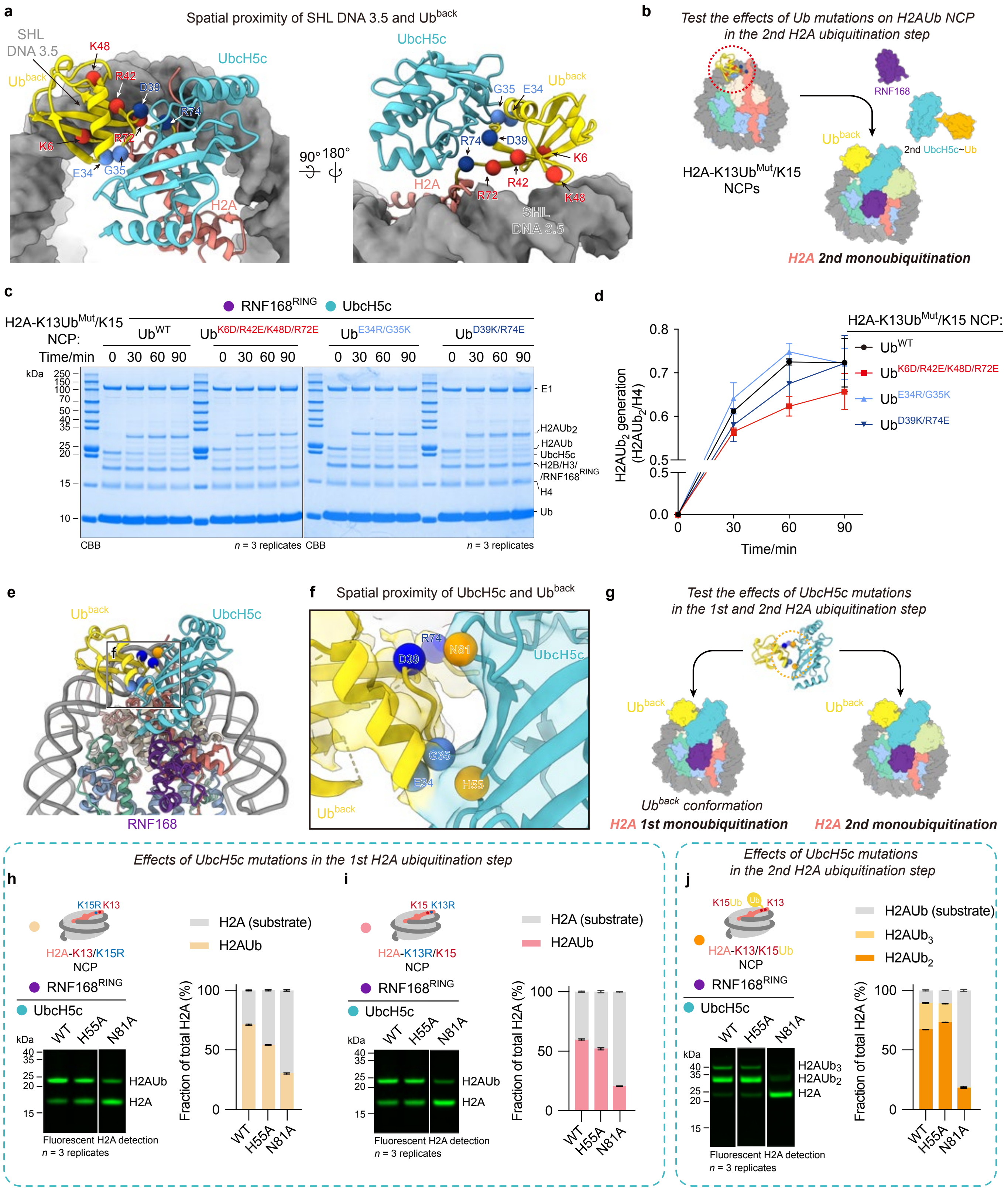


**Extended Data Figure 13. Biochemical investigations of the role of the structurally observed Ub^back in the RNF168-mediated H2A ubiquitinations.**

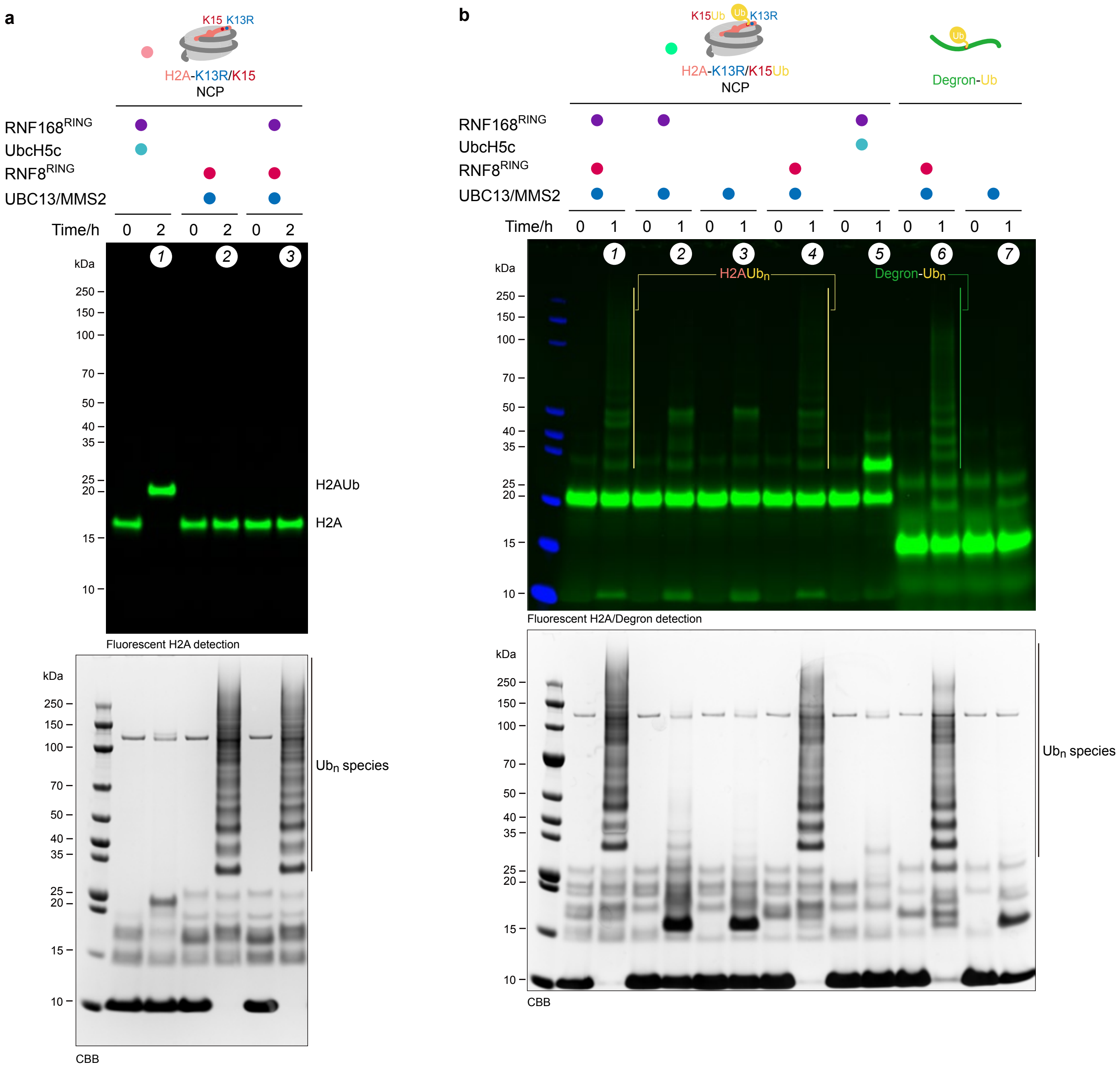


**Extended Data Figure 14. Biochemical investigations of roles of E3 ligase RNF8 in the RNF168-mediated H2A ubiquitinations.**
